# Repeated global migrations on different plant hosts by the tropical pathogen *Phytophthora palmivora*

**DOI:** 10.1101/2020.05.13.093211

**Authors:** Jianan Wang, Michael D. Coffey, Nicola De Maio, Erica M. Goss

**Affiliations:** Department of Plant Pathology and Emerging Pathogens Institute, University of Florida, Gainesville, FL, USA 32611; Department of Plant Pathology and Microbiology, University of California, Riverside, CA, USA 92521; Nuffield Department of Medicine, University of Oxford, OX3 7BN, Oxford, United Kingdom; European Molecular Biology Laboratory, European Bioinformatics Institute (EMBL-EBI), Wellcome Genome Campus, Hinxton CB10 1SD, UK

**Keywords:** host tracking, host shift, Oomycete, genetic diversity, structured coalescent analysis

## Abstract

The genetic structure and diversity of plant pathogen populations are the outcomes of evolutionary interactions with hosts and local environments, and migration at different scales, including human-enabled long-distance dispersal events. As a result, patterns of genetic variation in present populations may elucidate the history of pathogens. *Phytophthora palmivora* is a devastating oomycete that causes disease in a broad range of plant hosts in the tropics and subtropics worldwide. The center of diversity of *P. palmivora* is in Southeast Asia, but it is a destructive pathogen of hosts native to South America. Our objective was to use multilocus sequence analysis to resolve the origin and historical migration pathways of *P. palmivora*. Our analysis supports Southeast Asia as a center of diversity of *P. palmivora* and indicates that a single colonization event was responsible for the global pandemic of black pod disease of cacao. Analysis using the structured coalescent indicated that *P. palmivora* emerged on cacao and that cacao has been the major source of migrants to populations in Asia, Africa, and Pacific Islands. To explain these results, we hypothesize widespread introgression between the pandemic cacao lineage and populations native to Asia and the Pacific Islands. The complex evolutionary history of *P. palmivora* is a consequence of geographic isolation followed by long-distance movement and host jumps that allowed global expansion with cacao, coconut and other hosts.

---

With continued growth of human populations and change in global climate, there is increasing concern over plant diseases affecting food crop and economic security^1^. Epidemics of plant disease can result in major yield losses and associated economic consequences^2^. In low- and middle-income countries, plant diseases can have a relatively larger impact on local socio-economic development^3,4^. Understanding the population biology and evolution of plant pathogens can aid in disease management. For example, determining the geographic history of a plant pathogen can help locate plant germplasm that co-evolved with the pathogen and exhibits disease resistance, which can be employed in plant breeding programs. Knowledge of plant pathogen migration patterns can lead to informed decision-making for disease prevention, specifically to limit re-introductions and pathogen re-emergence^5^. Global pandemics may originate from a very successful invasive population that itself is responsible for multiple secondary invasions^3,6–8^. Hence, mitigation strategies may specifically target invasive populations or genotypes^9,10^.

However, the above efforts can be hampered by pathogens making dramatic host jumps. Host jumps are pervasive in the emergence or spread of plant pathogens and have been identified as a crucial mechanism underlying pathogen diversification and ultimately speciation^11,12^. International travel and global trade over the course of human history have exposed hosts to new pathogens, thus, facilitating host jumps^1,3,9,13–15^. Deciphering complex interactions among plant hosts, pathogens and human activities can elucidate major drivers behind the emergence, re-emergence and dispersal of plant pathogens^1^. Many such studies of pathogen emergence have targeted major crop pathogens that exhibit relatively narrow host associations^5,16–21^. When pathogens have multiple host associations, can the influence of different hosts on pathogen emergence and dispersal be resolved?

*Phytophthora* is a genus of plant-damaging Oomycetes, including more than 100 described species^22–25^. *Phytophthora* species can infect a broad range of plant hosts^26^ and have caused enormous economic losses to agro-ecosystems and ecological damage in natural ecosystems. The relatively narrow host range pathogen *Phytophthora infestans,* causal agent of potato and tomato late blight, was globally distributed on potato, leading to the Great Irish Famine. In 2008, *P. infestans* spread across the Eastern United States on tomato plants for home gardens^27^. *P. ramorum* is a broad host range pathogen, which dispersed on ornamental plants followed by host jumps to timber and forest trees^28^. The resulting disease outbreaks in North America and Europe have been economically costly and have changed the ecology of coastal California forests^29^. *Phytophthora palmivora* (Butler) Butler (1919) is a destructive tropical and subtropical plant pathogen with a very broad host range in the tropics and subtropics^30,31^*. P. palmivora* causes problematic diseases of coconut and other palms (bud rot), cacao (black pod, canker, and cherelle wilt), rubber (black stripe), durian (fruit rot and canker), orchids and other ornamentals, and more^26^. Annual losses due to black pod of cacao, caused primarily by *P. palmivora*, have been estimated at more than US$400 million per year^103^ and bud rot endemics have affected more than 70,000 ha of oil palm in Colombia^32^. However, like many other tropical pathogens, research on *P. palmivora* population biology and genetics has lagged far behind temperate *Phytophthora*, even though this pathogen is responsible for significant economic losses. To advance research on *P. palmivora*, the genome of an isolate from cacao was recently sequenced. Estimated genome size was greater than 151.23 Mb with 44 327 genes and initial analysis of gene models indicated that *P. palmivora* experienced a genome doubling event^33^.

The hypothesized center of origin of *P. palmivora* has changed over time. Initially, Central or South America was suspected to be the native region of the pathogen, because of the susceptibility of indigenous hosts and the apparent global distribution of the pathogen on cacao (*Theobroma cacao*), which is native to South America^34^. Findings of high levels of genetic diversity in *P. palmivora* populations in Southeast Asia, particularly among isolates from native hosts coconut and durian, led to the proposal that Southeast Asia is the center of origin of *P. palmivora*^30,35^. Recent advances in population genetics and coalescent model-based approaches have revolutionized methods to identify plant pathogen centers of origin, reconstruct migration pathways and reveal population genetic structure^36–44^. These new tools have substantially advanced knowledge of the evolutionary history and epidemiology of major plant pathogens, but have not been applied towards understanding the global spread of *P. palmivora*.

The main objective of this study was to describe the global population structure of *P. palmivora* and historical migration pathways using Bayesian and coalescent model-based inference approaches. We specifically addressed the following questions: (1) Are populations of *P. palmivora* structured by host or geography? (2) Where is the center of origin of *P. palmivora*? (3) What were the main migration pathways out of the center of origin? (4) Did host shifts drive the global expansion of the *P. palmivora*?

## Results

### Nucleotide diversity by geographic region

We evaluated DNA sequence variation in four genes (one mitochondrial and three nuclear) for three major geographic regions (Table 1). Higher values of average pairwise nucleotide diversity (π) and Watterson’s theta (θ_W_) were observed in the Asia-Pacific region than in Africa and the Americas. Tajima’s D and Fu & Li’s D* and F* test statistics were not significantly different from zero except for the sample from the Americas (Table 1). A significantly positive value of Tajima’s D was observed for *trp1* in the Americas, indicating more intermediate frequency alleles than expected under neutral evolution, while all three test statistics were significantly negative for *coxII* in the Americas, indicating excessive rare polymorphisms.

**Table 1.**
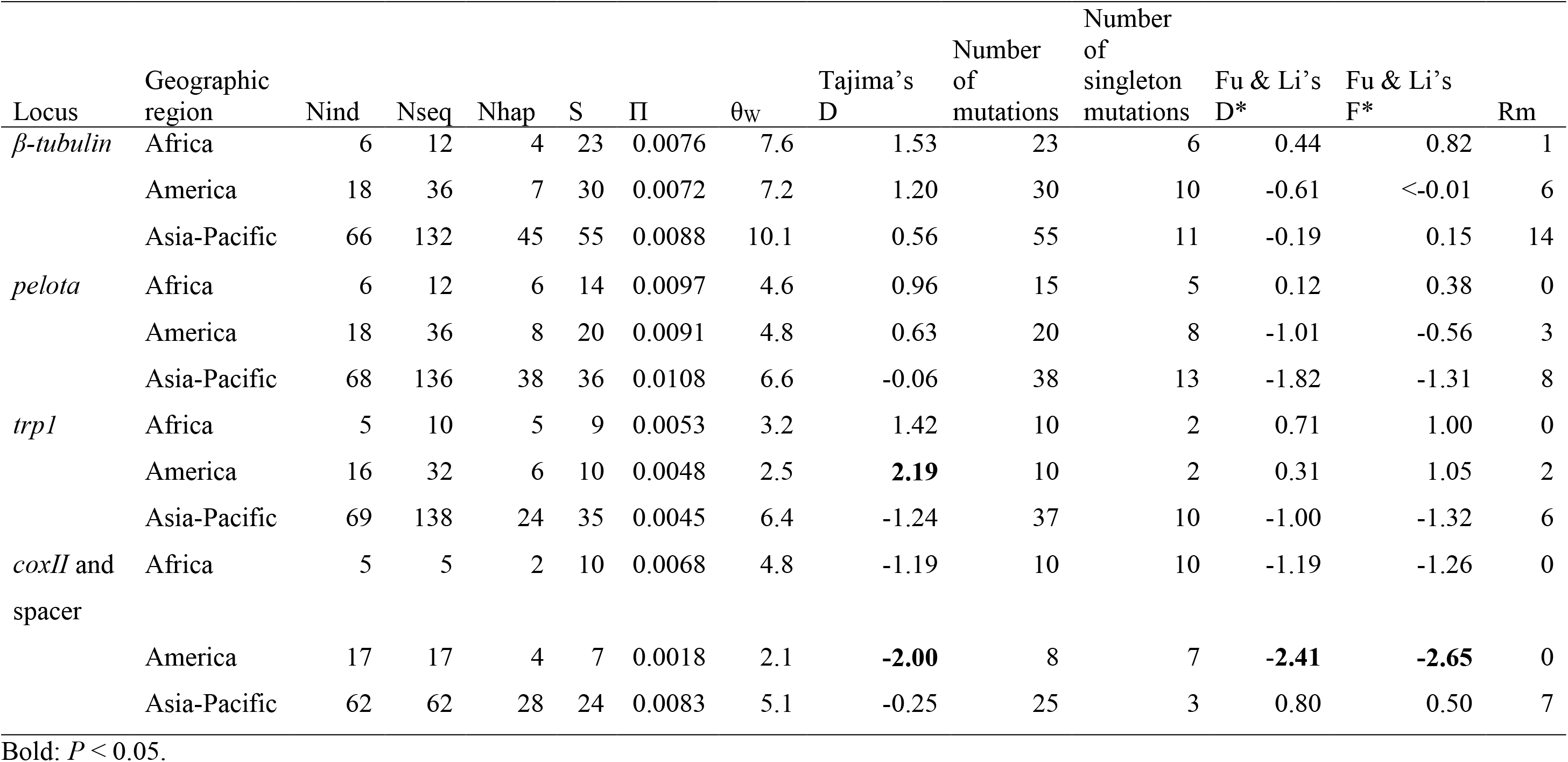
Nucleotide variation by locus and geographic region. Statistics given are number of individuals (Nind), number of sequences (Nseq), number of haplotypes (Nhap), segregating sites (S), average pairwise nucleotide diversity (π), Watterson’s theta (θW), Tajima’s D, number of mutations, number of singleton mutations, Fu and Li’s D* and F*, and minimum number of recombination events (Rm).

### Population subdivision and structure

Population differentiation was tested by AMOVA (Table 2). All four loci showed a significant portion of genetic variation (8-12%) distributed among regions (Africa, Americas and Asia-Pacific). Differentiation between the two most common hosts in the data set, coconut and cacao, explained around one quarter of the genetic variation in the nuclear loci (23-27%).

**Table 2.**
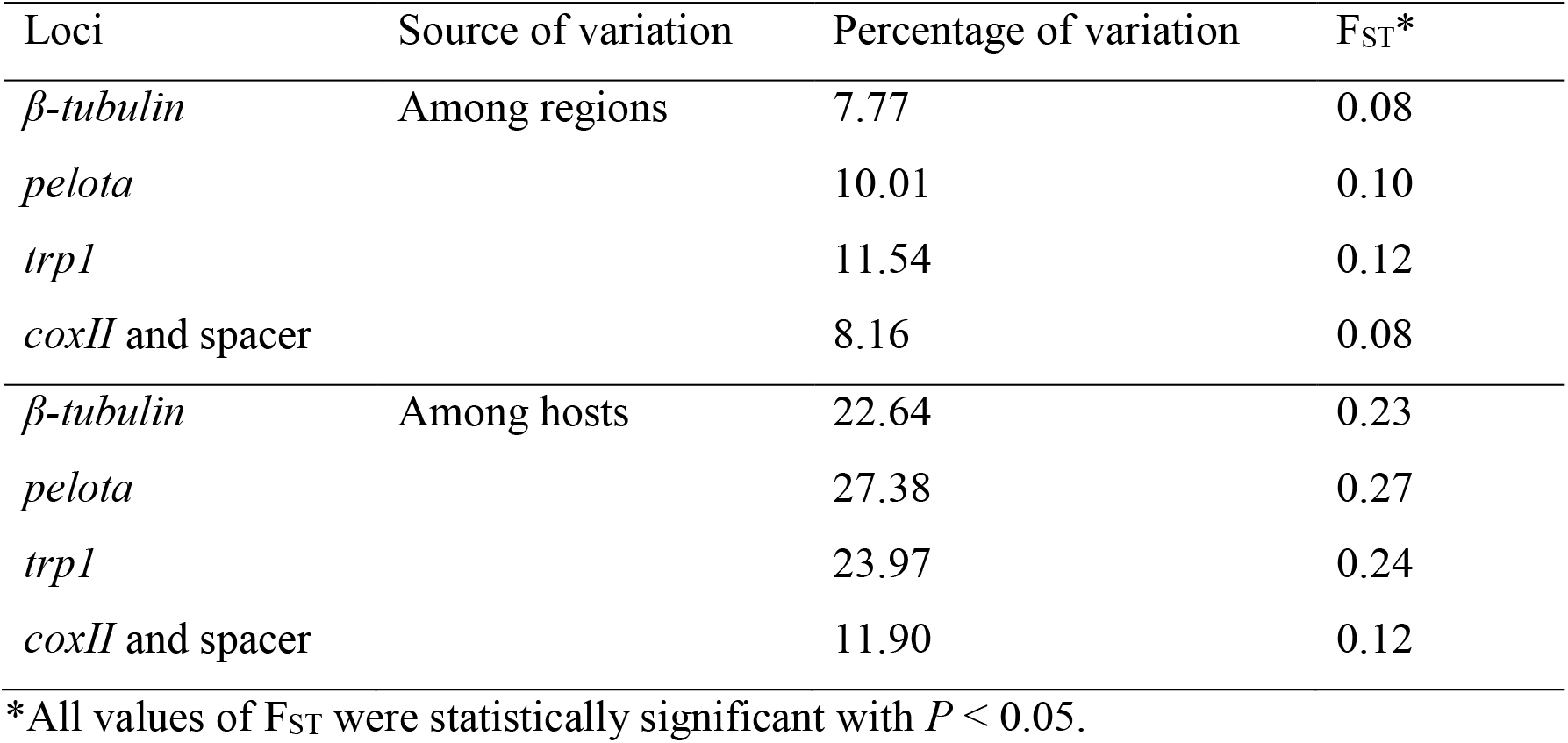
Analysis of molecular variance among three major geographic regions and two major hosts.

To further explore population structure, *P. palmivora* isolates were grouped by Bayesian clustering, using an allelic data set from the three nuclear loci (Fig. 1). When the number of clusters (K) was set to 2, most isolates from the Asia-Pacific region were in one cluster, and the majority of isolates from Africa and Americas were in the other. As K was increased, isolates from Indonesia, Philippines, Malaysia, and Pacific Island formed new clusters. Based on delta K, the optimum value of K was 2. The genetic structure within the Asia-Pacific region was greater than in Africa and the Americas. The small sample from south Asia (India and Sri Lanka) resembled populations in Africa and the Americas, the majority of which were isolated from cacao. Bayesian clustering of clone-corrected data produced the same overall pattern as the full data set (Supplementary Fig. 1).

**Fig. 1.**
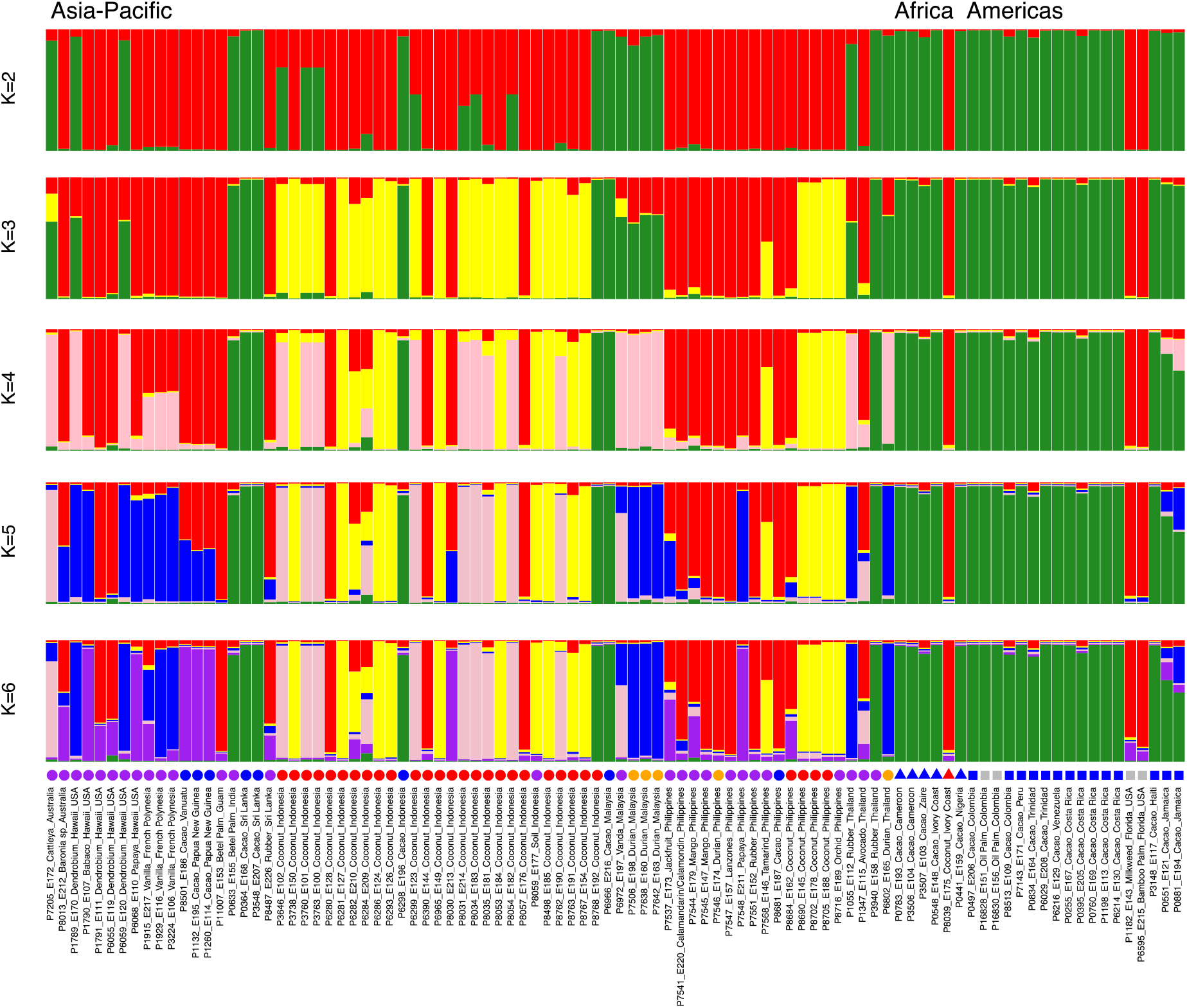
STRUCTURE assignments of *P. palmivora* isolates to populations for values of K from 2 to 6. Greater genetic structure is apparent in the Asia-Pacific region. Origins of isolates are indicated below each bar. Isolates from cacao are indicated by a blue symbol, those from coconut with a red symbol, and those from durian with an orange symbol. Isolates with purple or gray symbols are from other hosts. Region is indicated by the shape of the symbol (circle for Asia-Pacific, triangle for Africa, square for Americans).

The non-parameterized clustering method DAPC showed the first discriminant component separating isolates collected from cacao in all geographic regions from isolates collected from coconut and other hosts (Fig. 2). The second discriminant component separated isolates sampled from durian and other hosts from coconut and cacao isolates. Six isolates collected from coconut and other hosts in the Asia-Pacific and Americas clustered with isolates from cacao, consistent with isolates shifting from cacao to other hosts.

**Fig. 2.**
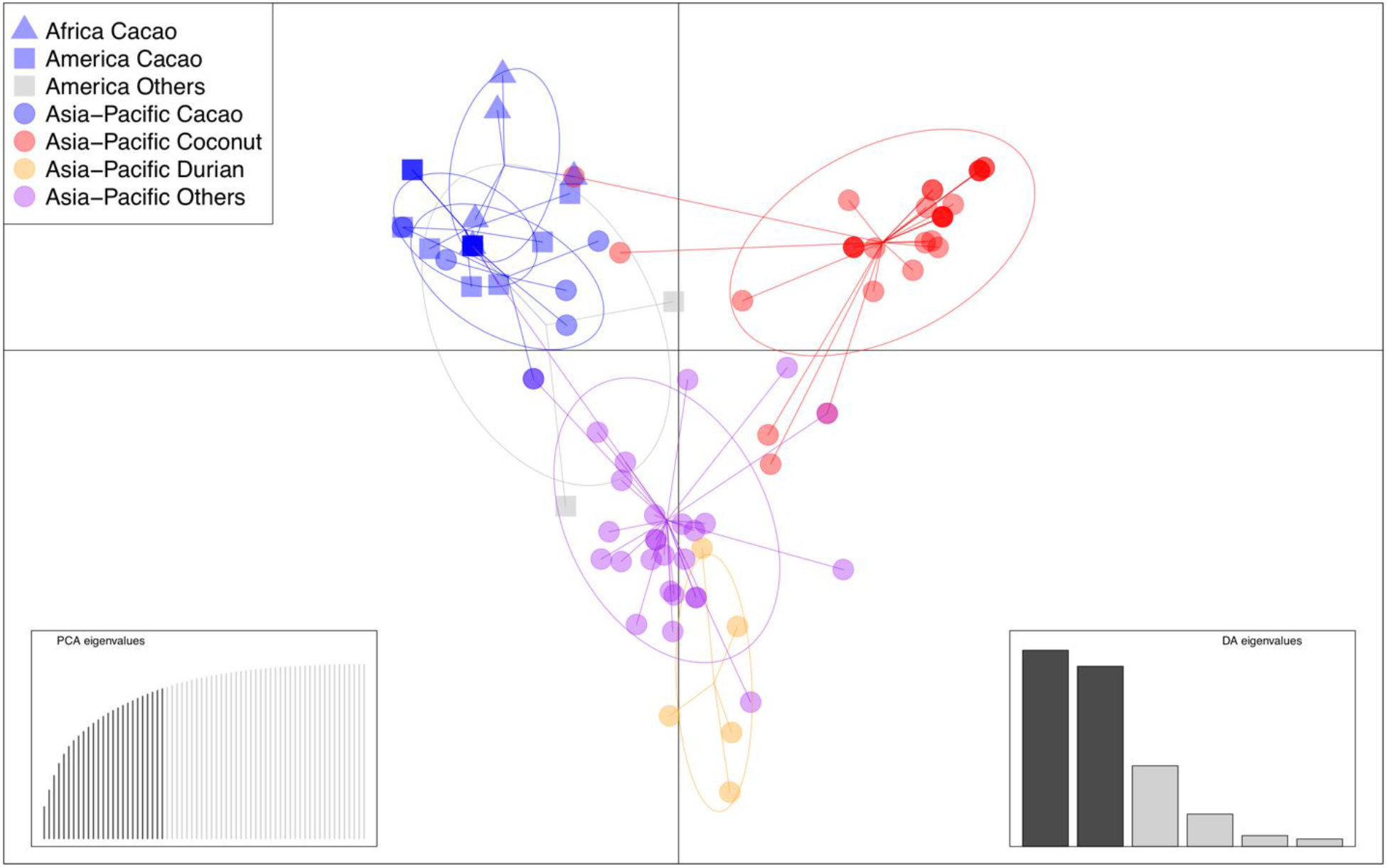
DAPC clusters of *P. palmivora* isolates from different hosts and geographic regions. All isolates collected from cacao in the three geographic regions grouped together. Asia-Pacific isolates sampled from coconut and durian grouped into two largely distinct clusters.

### Global migration patterns

Discrete Bayesian phylogeographic analysis was used to infer the geographic location of the root state of each locus^45,46^. The reconstructed root states supported Indonesia and/or Philippines as the inferred evolutionary origin of our sample (Fig. 3). The posterior probabilities for Philippines or Indonesia as the root state varied by locus (Supplementary Table 4). The corresponding analysis with BASTA, which is not as biased by sampling patterns, produced similar maximum clade credibility genealogies but with low posterior probabilities for geographic root states.

**Fig. 3.**
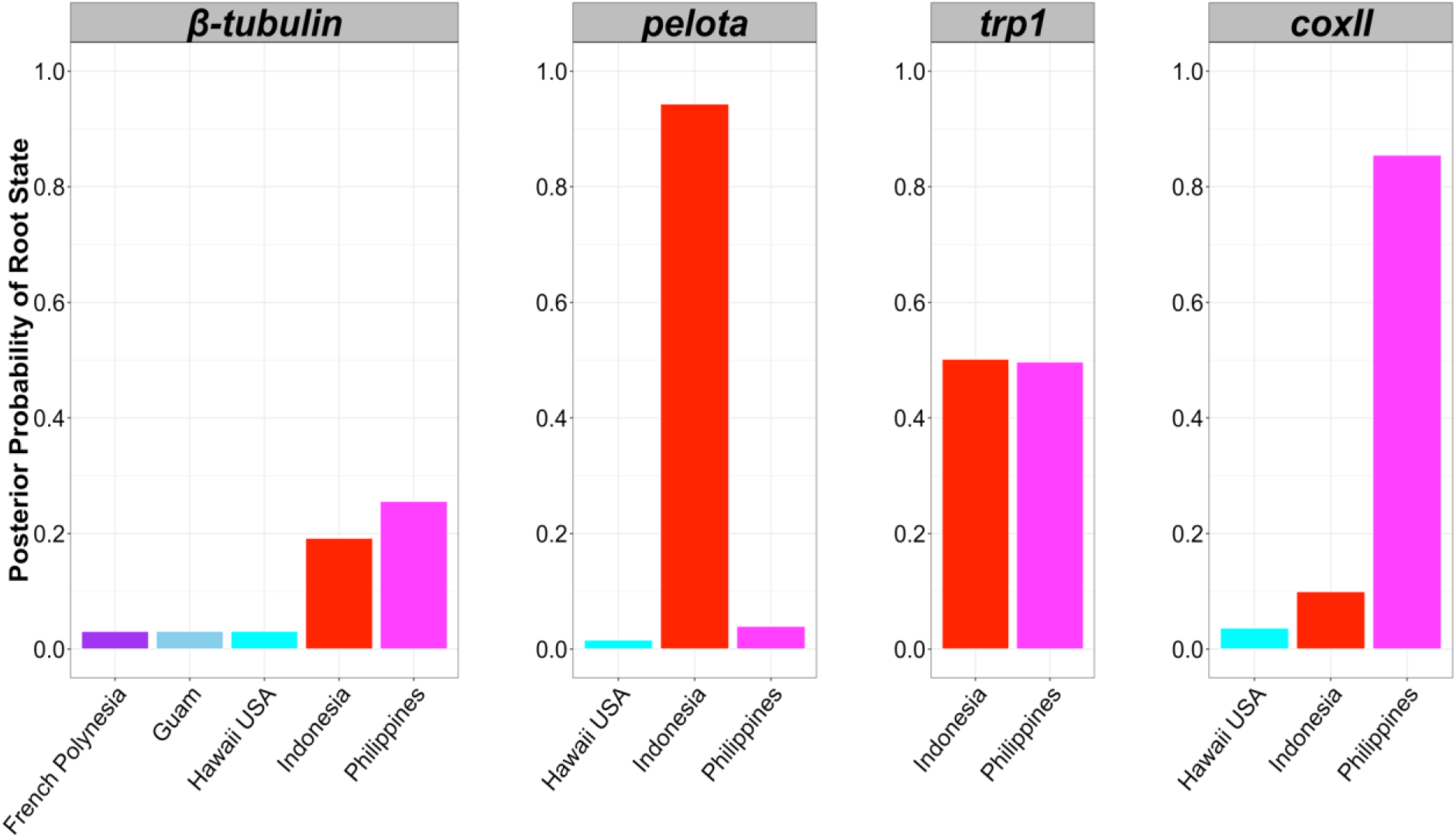
Root state posterior probability values inferred using discrete phylogeographic analysis for each location (country) by locus: *β-tubulin*, *pelota*, *trp1* and *coxII*. The posterior probabilities for an Indonesia root (red bar) and/or a Philippines root (magenta bar) are higher than for the other locations (other colors) for each locus, inferred independently. Due to limited space, only five locations with the five highest posterior probabilities for *β-tubulin* are shown. For *pelota*, *trp1* and *cox II*, only the locations with posterior probabilities more than 0.01 are shown.

The maximum clade credibility genealogies for the four loci (Fig. 4; Supplementary Figs. 2-4) indicate that lineages presently associated with Southeast Asia and the Pacific Islands diverged much earlier than lineages associated with Central and South America and Central Africa (mainly isolated from cacao). The mitochondrial locus, *coxII*, produced three lineages, one of which shows early diversification and is represented by isolates from SE Asia, Australia, Papua New Guinea, Vanuatu, French Polynesia, Hawaii, Ivory Coast and Jamaica (Fig. 4).

**Fig. 4.**
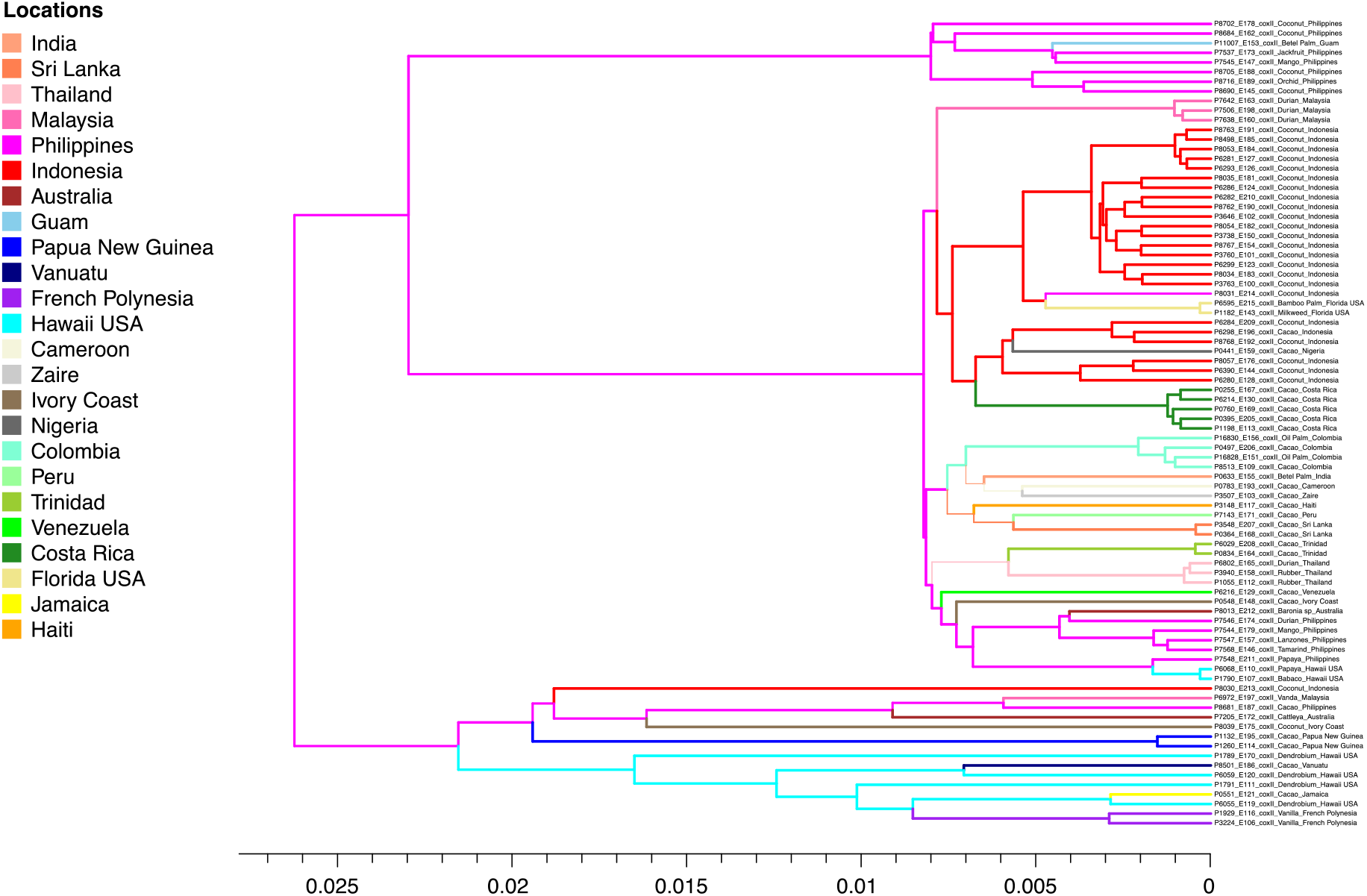
Maximum clade credibility genealogy for the *coxII* locus. Colors of branches indicate the most probable geographic origin of each lineage.

Across loci, isolates from Pacific Islands represented lineages that diverged prior to diversification of cacao-associated lineages. We observed that isolates from different hosts within given geographic regions had distinct evolutionary histories. For example, an isolate collected from coconut in Ivory Coast had distinct ancestry from the cacao isolates collected in Central Africa, and this was also observed for isolates from Florida and Jamaica. For the nuclear loci, we observed that most of the isolates from cacao exhibited two distinct haplotypes while showing little variation within each haplotype among isolates, consistent with a clonal lineage. Isolates from Indonesia and Philippines were less likely to have two diverged alleles but they exhibited more variation among isolates.

### Host switching

To complement the phylogeographic analysis and infer patterns of historical migration between major host groups, we conducted a multi-locus BASTA analysis by host group and region (Americas, Asia-Pacific and Africa). Based on analyses of genetic structure and phylogeography, we expected that migration occurred from coconut or other native hosts in SE Asia to hosts in the Pacific, to cacao in the Americas, and then back to cacao and other hosts in Asia. The BASTA analysis did not confirm this pattern. Using a two-host model to determine the directionality of gene flow between cacao and coconut, we inferred cacao to be the root host with high posterior probabilities (≥0.96 for each locus) and significant migration from cacao to coconut (Supplementary Table 5a). We examined if the existence of unsampled populations might have affected the analyses using a three-host model with one unsampled host. Cacao and the unsampled host were inferred as the root hosts with nearly even posterior probabilities. Mean rates of migration from cacao to coconut and the unsampled host to coconut were moderate to high, but the 95% HPD of the migration rates from the unsampled host included zero (Supplementary Table 5b), indicating uncertainty in these estimates. To incorporate geography, we examined a three-deme model, with populations cacao-America, cacao-Asia-Pacific and coconut-Asia. Cacao-America was inferred as the root state with high posterior probabilities (≥0.93 for each locus). Cacao-America was the major source of migrants, while cacao-Asia-Pacific and coconut-Asia were the major sinks (Table 3A; Supplementary Figs. 5-8). Incorporating an unsampled population into this model produced uncertainty in the root state with posterior probabilities of 0.63-0.68 for cacao-Asia-Pacific and 0.27-0.31 for the unsampled population and wide 95% HPDs on immigration estimates from the unsampled population (Table 3b). To explore if all remaining hosts, other than coconuts and cacao, in the Asia-Pacific region could represent the unsampled population, we examined a four-deme model replacing the unsampled deme with a putative deme we designated Other-Asia-Pacific. Similar to the previous models without an unsampled population, cacao-America was inferred as the root state with high posterior probabilities (≥0.96 for each locus), and cacao-Asia-Pacific and coconut-Asia were the two major sinks. Other-Asia-Pacific was inferred as a third major sink (Table 3c), which is the opposite migration direction to the models incorporating an unsampled deme. Models that explicitly incorporated the few cacao isolates from Africa produced results generally consistent with the models without the cacao-Africa deme (Supplementary Table 6a, b).

**Table 3.**
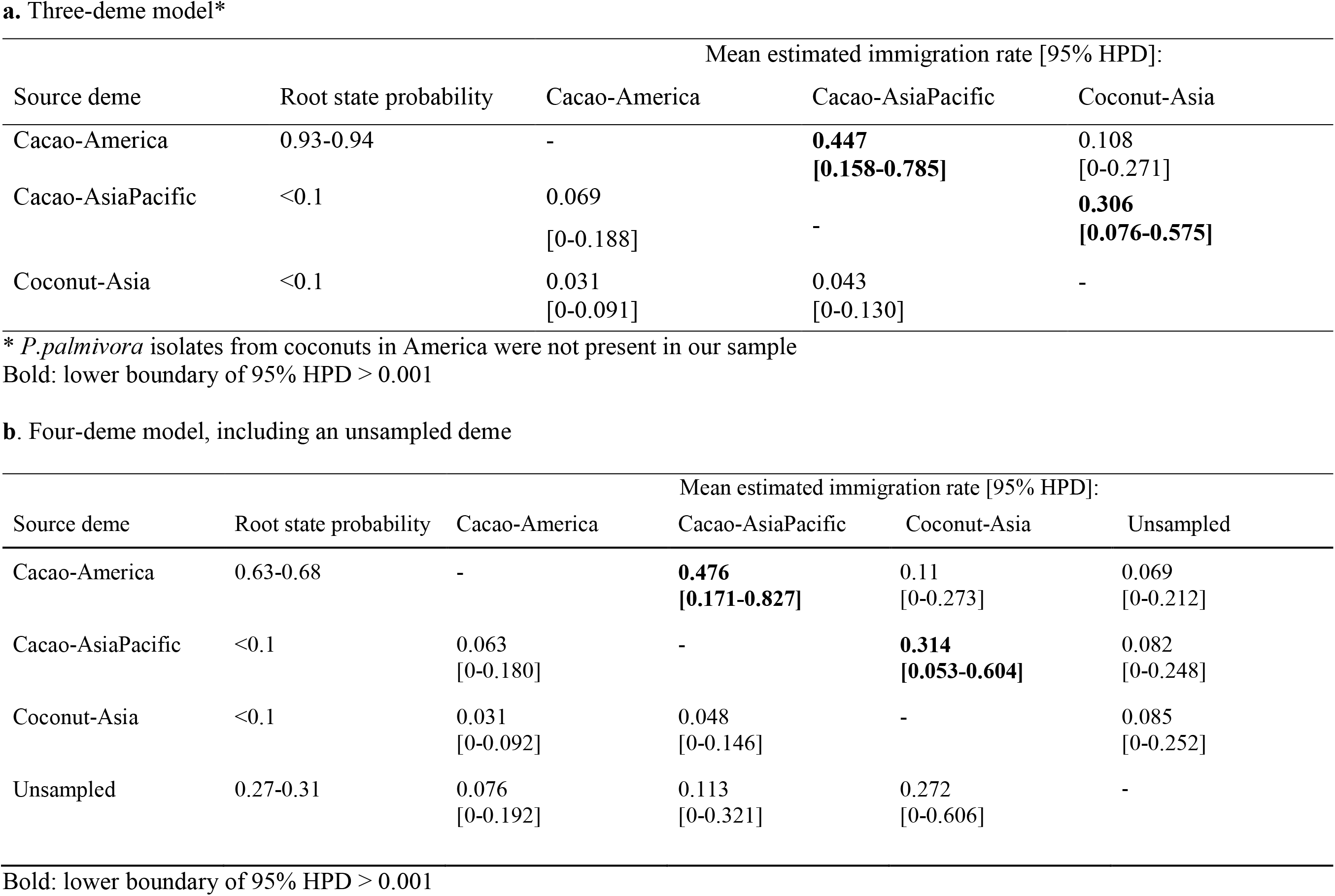
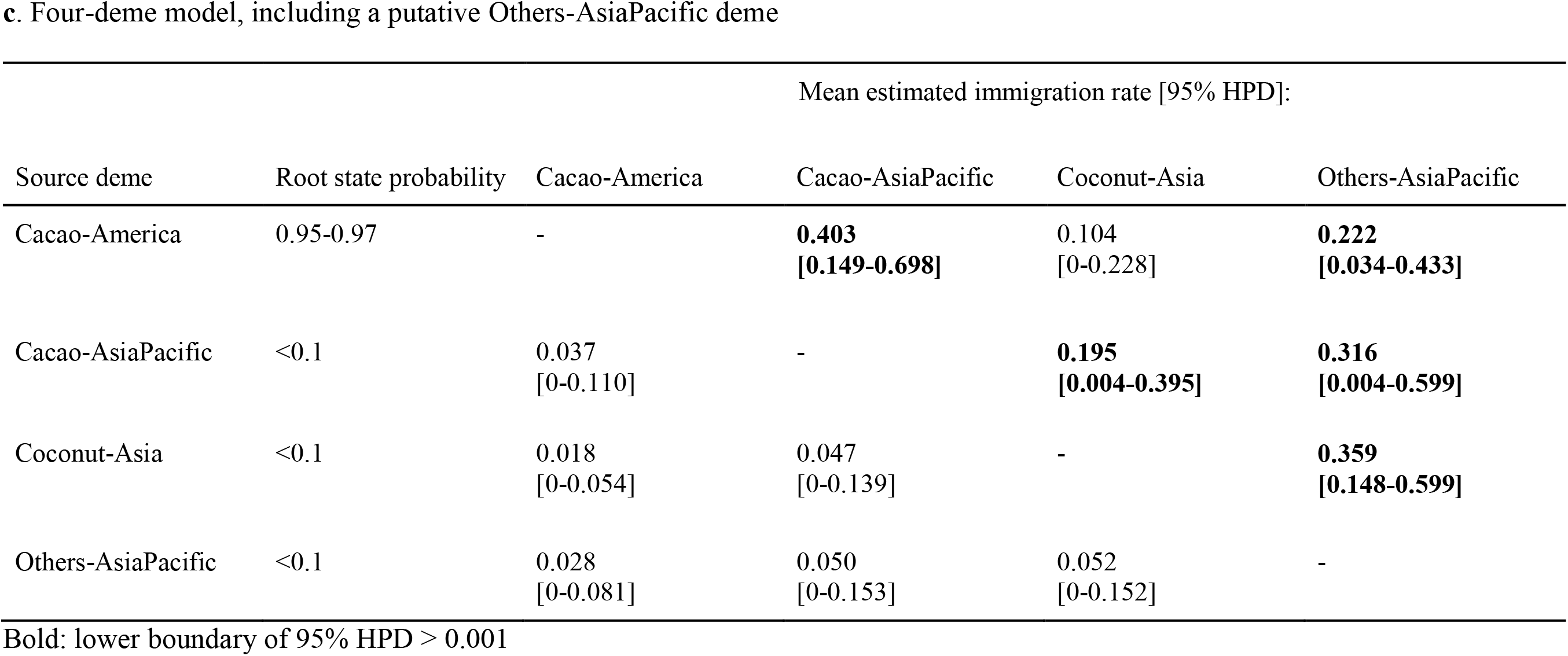
Estimates of migration rates between demes defined by hosts and geographical regions.

## Discussion

The evolutionary history of crop pathogens is as complex as the phylogeographic and agricultural histories of their hosts and the human populations that have cultivated them. Our multilocus sequence analysis of a global sample of *Phytophthora palmivora* isolates revealed population subdivision among three major geographic regions (Africa, Americas, Asia-Pacific) and between hosts, here highly sampled cacao and coconut. We found greater genetic diversity and more complex population structure within Southeast Asia as compared to other global regions, and reduced genetic variation among isolates collected from cacao. Therefore, our results generally support Southeast Asia as the *P. palmivora* center of diversity^30,35^. On the other hand, our analyses suggest that the Americas are a significant source of the genetic diversity now observed in the Asia-Pacific region. Here, we argue that the complex evolutionary history of *P. palmivora* reflects the history of human travel and trade in the tropics.

The greatest genetic diversity in *P. palmivora* appears to be within and among Pacific island chains, which naturally introduce geographic isolation due to historically infrequent movement of plant hosts, including by human migrations^47^. Genetically diverse isolates were associated with coconut palms and a variety of other tropical hosts. However, because the distribution of hosts sampled was not independent of geographic region, we cannot fully separate the relative influence of host and geography. For example, American and African isolates largely clustered together and separately from isolates from the Asia-Pacific region, but cacao was the most-sampled host in Africa and the Americas. Indeed, the population genetic structure of *P. palmivora* on cacao suggests that a single colonization was responsible for the global pandemic of *P. palmivora* causing black pod of cacao. The globally successful “cacao strain” is characterized by a single *coxII* haplotype and STRUCTURE cluster based on the genotypes of three nuclear loci. The most frequent haplotypes for each locus are associated with cacao. Many of the isolates from the Americas, Africa, and South Asia represented this cacao-associated lineage. The resulting pattern of genetic variation in the Americas produced positive values of Tajima’s D test across nuclear housekeeping genes and a statistically significant negative value for the mitochondrial *coxII* locus. These values are consistent with a recent population bottleneck^48^. In the Americas and South Asia, the “cacao strain” was isolated from palms and rubber, suggesting movement from cacao to other hosts. For example, two isolates obtained from cacao and oil palm in Colombia had identical sequences at the four sequenced loci. These isolates shared haplotypes with isolates from cacao worldwide and commercial oil palm production only began in Colombia in 1945^102^, therefore we infer that *P. palmivora* colonized oil palm from cacao.

We conducted phylogeographic analyses to infer the potential historical processes behind the observed patterns of genetic variation. Using discrete phylogeography, we inferred root states of our *P. palmivora* sample to be Indonesia or the Philippines, depending on the locus. We might expect that the pathogen has a metapopulation structure across these island nations, mediated by sea level changes joining or separating islands, dispersal of infected fruits following sea currents, or, more recently, trade in hosts infected by *P. palmivora*. The corresponding BASTA analysis was not able to infer a root location, and our reduced parameter analyses that grouped isolates by geographic region and host group inferred an American origin. BASTA is less susceptible to the effects of sampling bias, but inference of an American origin was unexpected because of the higher genetic diversity of the pathogen in the Asia-Pacific region. Simulations suggest that high posterior probabilities provided by the discrete model likely underestimate posterior uncertainty^44^. Our discrete model results reflect deep sampling in Southeast Asia, while the BASTA results indicate that sampling across a greater diversity of hosts and locations is needed. A long history of controversy regarding the geographic and evolutionary origin of the late blight pathogen, *P. infestans*, has been exacerbated by limited sampling of wild relatives of potato in the South American Andes^49,50^ and the geographic origins of many other widespread *Phytophthora* pathogens are unknown.

Migration estimates over the history of our sample of *P. palmivora*, inferred by BASTA, suggests movement from cacao in the Americas to cacao in Asia and Africa, and from cacao to other hosts in Asia. Because we observed genetic diversity on other hosts, it is unlikely that the sole origin of the pathogen is cacao. Indeed, we could not exclude the presence of an alternative, unsampled source population. In our inferred genealogies, we observed that haplotypes were not segregating independently, particularly for isolates from cacao, which likely represent an asexually reproducing lineage. The phylogeny for the *coxII* mitochondrial locus showed two groups of haplotypes, a large group of closely related haplotypes and a second group of low frequency, diversified haplotypes, which could represent different histories of ancestral populations of *P. palmivora*. One explanation for the observed patterns is introgression, perhaps repeated introgression, from the pandemic lineage on cacao to endemic populations of *Phytophthora* in Southeast Asia and the Pacific Islands. We do not yet have evidence of populations of *P. palmivora* or a closely related species on wild plants in the Americas which could have sourced migration to Asia. However, *P. palmivora* infects important tropical cash crops native to South America, including cacao^52^, rubber^53^, and neotropical orchids^54^. Genome sequencing of an isolate of *P. palmivora* from cacao indicates that this pathogen underwent genome doubling. We speculate that genome doubling was the consequence of a hybridization event between strains of different geographic origin, because genome doubling is commonly associated with interspecific hybridization^51^.

We can draw parallels between global patterns in the genetic variation of *P. palmivora* and historical global movement of the hosts considered here (Fig. 5). The Philippines represents one of two independent domestications of coconut (*Cocos nucifera*)^47,55^. Historical and genetic data have clarified the routes of coconut as it spread throughout the tropics and subtropics^55^. Various lines of evidence suggest that Pacific coconut was brought from the Philippines to eastern Polynesia and to the Pacific coast of Latin America by pre-Columbian Austronesians. Trade occurred in the opposite direction as well, because herbarium specimens and anthropological studies suggest pre-historic introduction of sweet potato from the Pacific coast of South America to Polynesia^56^. Later, the Portuguese set up plantations of the South Asian domestication (Indo-Atlantic genotypes) of coconut in West Africa, Brazil, and the Caribbean. Our findings of early divergence in our *P. palmivora* sample from the Pacific Islands is consistent with early movement, possibly on Pacific coconut, whereas isolates from Ivory Coast and Jamaica with deep nodes suggest movement on Indo-Atlantic coconut. *P. palmivora* infects numerous other tropical fruits and ornamentals native to the Asia-Pacific region, on which the pathogen could have dispersed or persisted over hundreds of years. If we assume that the divergence in sequences of isolates from Hawaii and other Pacific Islands are associated with human colonization of the Eastern Pacific islands around 1200 AD, we find that the colonization of global cacao production is during the last 150-200 years, as expected. The major documented movements of cacao to Africa and Asia was during the colonial period, and originated from Venezuela, Trinidad and Brazil^52,57^. Rubber is another major South American export and host of *P. palmivora*, with modern plantations located in South and Southeast Asia and West Africa^59,60^. The history of rubber in Asia started with the introduction of specimen trees from the Amazon region of Brazil to Ceylon, Singapore, and Penang in 1876, and rubber was established in plantations in Malaysia through 1898^58^.

**Fig. 5.**
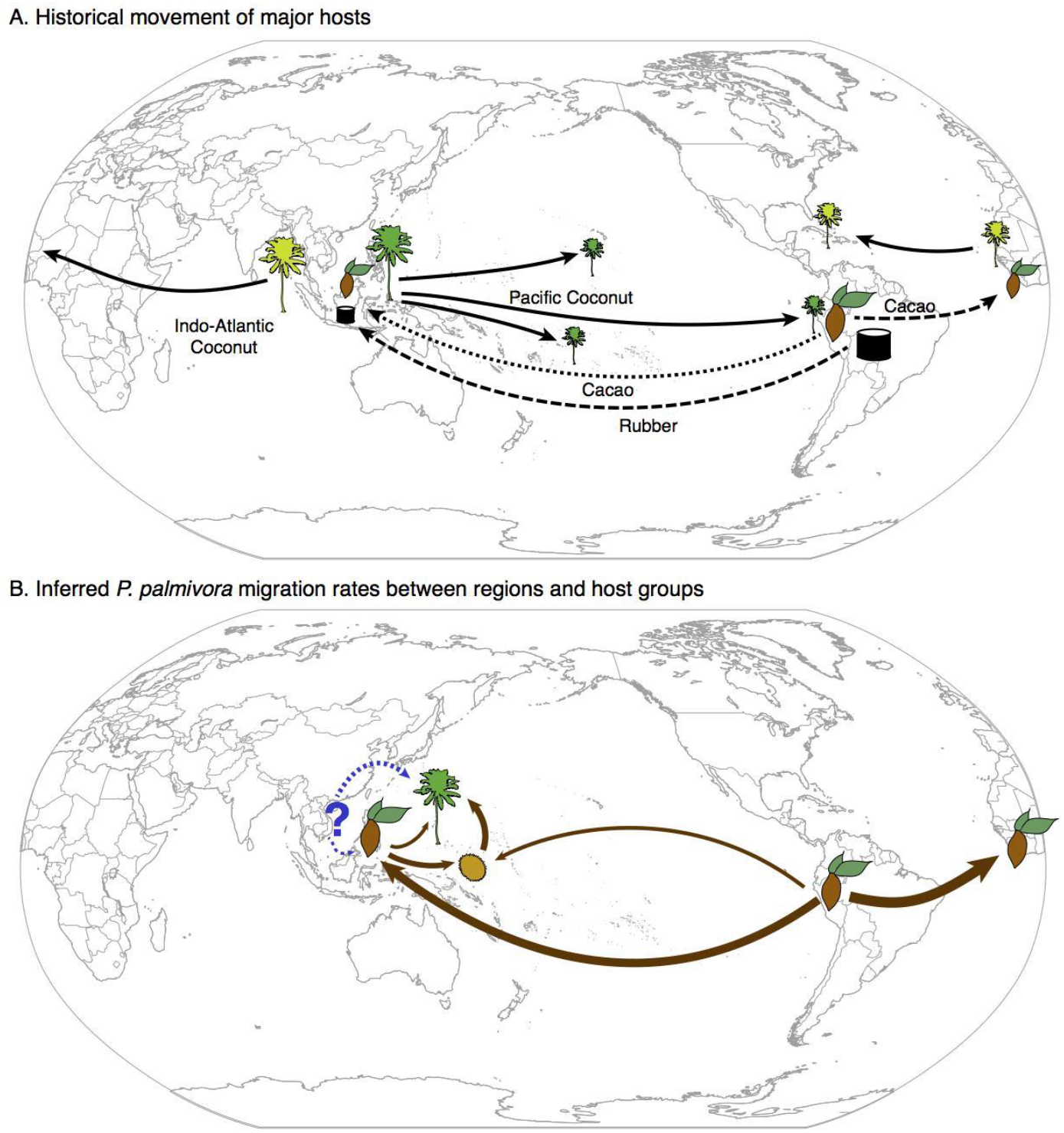
Movement of major hosts of *P. palmivora* out of their centers of origin and inferred migration of *P. palmivora* by host group. (**a**) Simplified historical movement of major *P. palmivora* hosts. Indo-Atlantic and Pacific coconut were domesticated in South and Southeast Asia, respectively. Cacao and rubber originated in South America. Many other tropical crops host *P. palmivora*, including ornamentals. (**b**) Summary of BASTA inferences of *P. palmivora* migration (solid lines, width proportional to mean migration rate) and potential migration from an unsampled deme (dotted lines). BASTA inferred high migration rates from cacao in the Americas to cacao in Asia-Pacific and Africa, and migration from cacao in Asia-Pacific to coconut and other hosts (indicated by a durian fruit), as well as migration from cacao in the Americas directly to other Asia-Pacific hosts.

We hypothesize that the diversity and genetic divergence of *P. palmivora* isolates in the Asia-Pacific region is an example of host-tracking. Our suggest that there may a long history of association of *P. palmivora* with coconut. Alternatively, *P. palmivora* may have long-term associations with other hosts native to Southeast Asia and coconut served primarily as an early vector for long-distance dispersal. The host-tracking model of plant pathogen emergence has been seen in other host-pathogen systems in Southeast Asia, such as the black Sigatoka pathogen *Mycosphaerella fijiensis* on banana^61^ and blast fungus *Magnaporthe oryzae* on rice^62^. In contrast, the nearly monomorphic population *P. palmivora* found in Americas, South Asia, and Africa can be explained by a pandemic lineage of *P. palmivora* on cacao, following a genetic bottleneck likely associated with initial colonization of this host. The population of *P. palmivora* associated with cacao resembles the pattern of emergence of globally-distributed clonal lineages of *P. infestans* on potato^21,63–65^. Several other oomycetes have exhibited similar genetic bottlenecks upon intercontintental spread, including *Phytophthora ramorum*^28^. These events are often associated with a host jump, particularly colonization of plants for export.

The evolutionary and geographic history of *P. palmivora* could be clarified by examination of additional isolates from uncultivated plants, which may represent older or more isolated populations, or by using preserved DNA from herbarium samples. In general, the distributions of crop pests and pathogens are highly dependent on the distribution of their hosts, even though the specific mechanisms of global spread differ among species^66^. The idiosyncrasies of plant movement during human history shaped the genetic variation and structure of modern populations of cultivated plants and of their pathogens. Therefore, efforts to manage disease may benefit from teasing apart historical dependencies that structure host and pathogen genetic variation and that are likely to mediate the plant-pathogen interaction.

## Methods

### *P. palmivora* isolates

Genomic DNA was provided from 95 isolates of *P. palmivora* in the World Oomycete Genetic Resource Collection at the University of California, Riverside, USA. The isolates were originally collected from 23 hosts and 22 countries on five continents (Africa, Asia, Australia, North America and South America) (Supplementary Table 1). For analysis, isolates were assigned to one of three large geographical regions: 1) Asia-Pacific (Asia, Australia and Pacific Islands); 2) Americas (including the Caribbean islands); and 3) Africa.

### Multilocus sequence typing

Four housekeeping genes known to contain variation within *Phytophthora* species were used in this study: mitochondrial gene *coxII* with the adjacent spacer^67^, and three nuclear genes, *β-tubulin*, *pelota* and *trp1*^68,69^. Primers for the nuclear genes were modified for *P. palmivora* based on draft whole genome sequence kindly provided by S. Schornack and S. Kamoun (Supplementary Table 2).

For PCR, a total reaction volume of 25.0 μl was prepared including 1.0 ul DNA template, 1X OneTaq master mix (NEB), 0.2uM primers and 3.0 uM MgCl_2_. PCR amplifications were performed using the following cycling protocol: initial denaturation at 94 °C for 3 minutes; followed by 35 cycles of 94 °C for 45 seconds, [(67 °C for 45 seconds (*pelota*); 58 °C for 45 seconds (*β-tubulin*); 60 °C for 45 seconds (*coxII*); 59°C for 60 seconds (*trp1*)) and 68 °C for 1 minute; then a final extension step was 68 °C for 10 minutes. PCR products were prepared for sequencing using ExoSAP-IT (USB Corporation, Cleveland, OH, USA) and directly sequenced in both directions at Interdisciplinary Center for Biotechnology Research (ICBR) at the University of Florida, USA. Reads were assembled and edited using software Geneious 6.1.8 (https://www.geneious.com). For the three nuclear loci, haplotype phase was inferred using the program PHASE^70^, assuming diploidy. The settings for PHASE were as follows: MR0 (the default model, which is the general model for recombination rate variation^71^); number of iterations=10000 for *β-tubulin* and *trp1*, 2000 for *pelota*; thinning interval=1; burn-in=100. For each nuclear locus, ten isolates that exhibited more than one heterozygous site were randomly selected and cloned using the pGEM-T Easy Vector System (Promega Corporation, Madison, WI, USA). Sequences were aligned with Geneious 6.1.8^72^. Indels were removed for analysis.

### Nucleotide diversity and neutrality tests

Number of individuals (Nind), number of sequences (Nseq), number of haplotypes (Nhap), segregating sites (S), average pairwise nucleotide diversity (π)^73,74^, Watterson’s theta (θ_w_)^75^, Tajima’s D^76–79^, number of mutations, number of singleton mutations, Fu and Li’s D* and F*^76,77,80–85^ and minimum number of recombination events (Rm)^86^ were determined for each of the four genes and by region using DnaSP v5^87^.

### Population subdivision and structure

Population subdivision among the three major geographic regions (Africa, Americas and Asia-Pacific) and between two major hosts (coconut, native to Southeast Asia; and cacao, native to South America) were examined for each gene locus by analysis of molecular variance (AMOVA) in Arlequin 3.5.2.1^88^.

The population structure of *P. palmivora* was examined by model-based Bayesian clustering carried out in STRUCTURE 2.3.4^36^. For this analysis, allelic data were used from the three nuclear loci. The program STRUCTURE estimates parameters independently in the posterior probability distribution of allele frequencies. Parameters are estimated under the null model of panmixia within populations, which is characterized by Hardy-Weinberg equilibrium at each locus and an absence of linkage disequilibrium. The algorithm is robust to some departures from these assumptions^89–91^. Based on allele frequencies at each locus, STRUCTURE assigns Q-values (inferred ancestry) to each individual. Isolate P3444 was excluded, due to missing data for *pelota* and *trp1*. To reduce the effect of asexual reproduction on STRUCTURE inferences, two different input files were prepared. One used the full data set of 94 isolates and a second used a clone-corrected data set of 69 isolates. Clone correction is a method to remove bias caused by dominant or overrepresented genotypes generated by epidemic asexual reproduction. Here, we clone-corrected globally by including one representative isolate of each multilocus genotype. We applied the following workflow to analyze both input files. STRUCTURE was run using the admixture model, and cluster numbers (K) from K=1 to K=15 were evaluated using 500 000 iterations after a burn-in period of 500 000 iterations. To evaluate the stability of the results across repeated runs, 20 independent runs were conducted. The runs for each value of K were evaluated based on the second order rate of change of the likelihood function with respect to K^92^ using the online program *Structure Harvester* v.0.6.94^93^. Due to stochastic effects among replicate STRUCTURE runs, assignment probabilities were compiled for multiple runs in the program CLUMPP version 1.2, which simplifies the assessment of replicate data by calculating medians^94^. The parameters used were M=3, W=0 and S=2, GREEDY_OPTION=2, and REPEATS=10000. The graphical visualization of the output was produced using R base function ‘*barplot’* ^95^.

Population subdivision, based on allelic data from the three nuclear genes, was also examined using model-free discriminant analysis of principal components (DAPC) implemented in the R package *adegenet* 1.4.2^41^. DAPC was run to confirm the results of Bayesian clustering approach. DAPC does not make biological assumptions and is less computationally intensive than STRUCTURE. This method maximizes the variation between groups while minimizing variation within group. First, DAPC transforms the data using principal components analysis (PCA) and then discriminant analysis (DA) assigns individuals to clusters. The data transformation ensures that the variables inputted to DA are uncorrelated and that their number is less than that of analyzed individuals, so as to overcome the drawbacks of direct application of DA. The appropriate number of principal components retained in the analysis can be determined by cross-validation^41^. Because only one isolate, P8039, was in the pre-defined coconut-Africa group and could not be used for in both training and validation sets, we removed this isolate and retained 93 isolates for the DAPC analyses.

### Genealogies and Phylogeographic Analysis

To infer the phylogeographic history of *P. palmivora*, genealogies were inferred by Bayesian evolutionary analysis by sampling trees (BEAST) and location states by discrete trait analysis^37,46^. We conducted separate analyses for each locus, each of which is expected to have a different genealogy from the others. BEAST executes ancestral reconstruction of discrete states, here location of collection, in a Bayesian statistical framework for evolutionary hypothesis testing using rooted, time-measured phylogenies. In this analysis, locations associated with branch nodes were characterized by continuous time Markov chain models, comparable to nucleotide, codon or amino acid substitution models and Bayesian stochastic search variable selection (BSSVS) was used to model the phylogeographic dispersion process^45^. This approach uses the geographic locations of the samples to reconstruct the ancestral states of tree nodes and the root state. A strict molecular clock model was used. The mutation rate was set as a constant 1.0, consequently the estimation of branch lengths is in substitutions per site. We used jModelTest 2.1^96^ to obtain the best fitting models of nucleotide substitution for each gene alignment. A coalescent tree prior and a prior of constant population size were assumed. Three replications of independent MCMC were run for each locus and the first 10% of recorded states were discarded as burn-in (parameter settings in Supplementary Table 3). The program was run until effective sample size estimates of greater than 200 were obtained, with good mixing and convergence in independent runs, as assessed in Tracer v1.6^97^. Maximum clade credibility (MCC) trees were summarized by TreeAnnotator based on common ancestor height^98^ and visualized using R package *rBt*^99^.

We complemented the above analysis with Bayesian structured coalescent approximation (BASTA)^44^ in BEAST 2.0^100^. BASTA incorporates migration into the structured coalescent-based model rather than modeling migration independently from the coalescent process as in the above discrete phylogeographic analysis. BASTA phylogeographic inferences are less influenced by sampling bias and variation in sampling intensity, and can include unsampled ghost demes. All four loci were used in the analysis, were assumed to be unlinked, and were assigned different nucleotide substitution models, as determined using jModeltest2 for each locus. The symmetric BSSVS with 24 locations (shown in Fig. 4) was used to model the phylogeographic dispersion process, which assumed a prior in which 60 migration rates are expected to be non-zero^45^. We assumed the same population size for all demes. Two billion steps were used in the MCMC chain for each run, and one tree was recorded every 200 000 steps. To reduce the lengthy running time and to obtain an effective sample size estimates of greater than 200 with good mixing and convergence, two parallel runs with different seed numbers were implemented, and the two parallel tree files for each locus were merged. The merged tree files were assessed in Tracer v1.6^97^. After the first 10% of recorded ‘burn-in’ states were discarded, MCC trees were summarized by TreeAnnotator based on common ancestor height^98^.

We also used BASTA to explore major between-host transmissions in relation to geography by estimating migration rates between demes defined by host and region. We started with a simple model of the two major hosts, cacao and coconut, to determine which of these two is more likely to be the ancestral host of *P. palmivora*. We incorporated geography by splitting cacao isolates into cacao-Asia-Pacific and cacao-America demes, resulting in three demes: cacao-America, cacao-Asia-Pacific and coconut-Asia. We also examined the effect of adding an unsampled (ghost) deme. Because we did not include isolates collected from hosts other than coconuts and cacao in the Asia-Pacific region, we grouped these isolates into a deme to determine if these hosts represented the unsampled (ghost) deme. Finally, we examined the effect on the above models of adding a cacao-Africa deme, although the sample size from Africa is small. The four loci were assumed to be unlinked and assigned different nucleotide substitution models, as above. We ran migration models that allowed for asymmetric migration between demes, assuming the same population size for all demes. Three hundred million steps were used in the MCMC chain for each run, and one tree was recorded every 100 000 steps. Mixing was assessed and trees examined as above. The MCC trees were visualized by IcyTree^101^. Two replications of independent MCMC were run for each model.

## Data availability

The DNA sequence data, inferred haplotypes and the analysis scripts that support the findings of this study will be made available from the Dryad repository upon submission of this preprint to another journal.

## Acknowledgements

We thank Jordan McLendon for assistance in the laboratory, Sebastian Schornack and Sophien Kamoun for sharing *P. palmivora* genome data for primer design, and Santiago Sánchez-Ramírez for assistance with *rBt*. This work was supported by the University of Florida Department of Plant Pathology and Emerging Pathogens Institute.

## Author contributions

JW and EMG designed the experiments. MDC contributed the DNA samples of *Phytophthora palmivora* from the World Oomycete Genetic Resource Collection. JW performed the data analyses with contributions from EMG and NDM. JW and EMG wrote the manuscript with contribution from NDM and MDC. All authors reviewed and approved the manuscript.

