## Supplementary for "Repeated global migrations on different plant hosts by the tropical pathogen *Phytophthora palmivora*" (from another version of this preprint: version 1, 2020-05-14)

**Running title:** Host tracking by a broad host range plant pathogen

**Supplementary Table 1.** List of 95 *P. palmivora* isolates used in the study.

| Accession Number | DNA Number | Region | Country | Year | Host |
|---|---|---|---|---|---|
| P3763 | E100 | Asia-Pacific | Indonesia | 1988 | Coconut |
| P3760 | E101 | Asia-Pacific | Indonesia | 1988 | Coconut |
| P3646 | E102 | Asia-Pacific | Indonesia | 1988 | Coconut |
| P3507 | E103 | Africa | Zaire | 1959 | Cacao |
| P3506 | E104 | Africa | Cameroon | 1959 | Cacao |
| P3224 | E106 | Asia-Pacific | French Polynesia | 1986 | Vanilla |
| P1790 | E107 | Asia-Pacific | Hawaii, USA | 1983 | Babaco |
| P8513 | E109 | America | Colombia | 1994 | Cacao |
| P6068 | E110 | Asia-Pacific | Hawaii, USA | 1989 | Papaya |
| P1791 | E111 | Asia-Pacific | Hawaii, USA | 1983 | Dendrobium |
| P1055 | E112 | Asia-Pacific | Thailand | 1976 | Rubber |
| P1198 | E113 | America | Costa Rica | 1980 | Cacao |
| P1260 | E114 | Asia-Pacific | Papua New Guinea | 1980 | Cacao |
| P1347 | E115 | Asia-Pacific | Thailand | - | Avocado |
| P1929 | E116 | Asia-Pacific | French Polynesia | 1986 | Vanilla |
| P3148 | E117 | America | Haiti | 1987 | Cacao |
| P3444 | E118 | Asia-Pacific | Australia | 1988 | sugar apple |
| P6055 | E119 | Asia-Pacific | Hawaii, USA | 1989 | Dendrobium |
| P6059 | E120 | Asia-Pacific | Hawaii, USA | 1989 | Dendrobium |
| P0551 | E121 | America | Jamaica | 1968 | Cacao |
| P6299 | E123 | Asia-Pacific | Indonesia | 1989 | Coconut |
| P6286 | E124 | Asia-Pacific | Indonesia | 1989 | Coconut |
| P6293 | E126 | Asia-Pacific | Indonesia | 1989 | Coconut |
| P6281 | E127 | Asia-Pacific | Indonesia | 1989 | Coconut |
| P6280 | E128 | Asia-Pacific | Indonesia | 1989 | Coconut |
| P6216 | E129 | America | Venezuela | - | Cacao |
| P6214 | E130 | America | Costa Rica | - | Cacao |

**Supplementary Table 1** Continued

| Accession Number | DNA Number | Region | Country | Year | Host |
|---|---|---|---|---|---|
| P1182 | E143 | America | Florida, USA | 1979 | Milkweed |
| P6390 | E144 | Asia-Pacific | Indonesia | 1989 | Coconut |
| P8690 | E145 | Asia-Pacific | Philippines | - | Coconut |
| P7568 | E146 | Asia-Pacific | Philippines | 1992 | Tamarind |
| P7545 | E147 | Asia-Pacific | Philippines | 1992 | Mango |
| P0548 | E148 | Africa | Ivory Coast | 1968 | Cacao |
| P6965 | E149 | Asia-Pacific | Indonesia | 1990 | Coconut |
| P3738 | E150 | Asia-Pacific | Indonesia | - | Coconut |
| P16828 | E151 | America | Colombia | 2009 | Oil Palm |
| P7551 | E152 | Asia-Pacific | Philippines | 1992 | Rubber |
| P11007 | E153 | Asia-Pacific | Guam | 2004 | Betel Palm |
| P8767 | E154 | Asia-Pacific | Indonesia | 1994 | Coconut |
| P0633 | E155 | Asia-Pacific | India | - | Betel Palm |
| P16830 | E156 | America | Colombia | 2009 | Oil Palm |
| P7547 | E157 | Asia-Pacific | Philippines | 1992 | Lanzones |
| P3940 | E158 | Asia-Pacific | Thailand | - | Rubber |
| P0441 | E159 | Africa | Nigeria | - | Cacao |
| P7638 | E160 | Asia-Pacific | Malaysia | 1992 | Durian |
| P8684 | E162 | Asia-Pacific | Philippines | 1994 | Coconut |
| P7642 | E163 | Asia-Pacific | Malaysia | 1992 | Durian |
| P0834 | E164 | America | Trinidad | - | Cacao |
| P6802 | E165 | Asia-Pacific | Thailand | 1989 | Durian |
| P0255 | E167 | America | Costa Rica | 1964 | Cacao |
| P0364 | E168 | Asia-Pacific | Sri Lanka | 1965 | Cacao |
| P0760 | E169 | America | Costa Rica | 1971 | Cacao |
| P1789 | E170 | Asia-Pacific | Hawaii, USA | 1986 | Dendrobium |
| P7143 | E171 | America | Peru | 1990 | Cacao |
| P7205 | E172 | Asia-Pacific | Australia | 1990 | Cattleya |

**Supplementary Table 1** Continued

| Accession Number | DNA Number | Region | Country | Year | Host |
|---|---|---|---|---|---|
| P7537 | E173 | Asia-Pacific | Philippines | 1992 | Jackfruit |
| P7546 | E174 | Asia-Pacific | Philippines | 1992 | Durian |
| P8039 | E175 | Africa | Ivory Coast | 1993 | Coconut |
| P8057 | E176 | Asia-Pacific | Indonesia | 1993 | Coconut |
| P8059 | E177 | Asia-Pacific | Indonesia | 1993 | Soil |
| P8702 | E178 | Asia-Pacific | Philippines | - | Coconut |
| P7544 | E179 | Asia-Pacific | Philippines | 1992 | Mango |
| P8035 | E181 | Asia-Pacific | Indonesia | 1993 | Coconut |
| P8054 | E182 | Asia-Pacific | Indonesia | 1993 | Coconut |
| P8034 | E183 | Asia-Pacific | Indonesia | 1993 | Coconut |
| P8053 | E184 | Asia-Pacific | Indonesia | 1993 | Coconut |
| P8498 | E185 | Asia-Pacific | Indonesia | 1994 | Coconut |
| P8501 | E186 | Asia-Pacific | Vanuatu | 1994 | Cacao |
| P8681 | E187 | Asia-Pacific | Philippines | - | Cacao |
| P8705 | E188 | Asia-Pacific | Philippines | - | Coconut |
| P8716 | E189 | Asia-Pacific | Philippines | - | Orchid |
| P8762 | E190 | Asia-Pacific | Indonesia | 1994 | Coconut |
| P8763 | E191 | Asia-Pacific | Indonesia | 1994 | Coconut |
| P8768 | E192 | Asia-Pacific | Indonesia | 1994 | Coconut |
| P0783 | E193 | Africa | Cameroon | 1971 | Cacao |
| P0881 | E194 | America | Jamaica | 1977 | Cacao |
| P1132 | E195 | Asia-Pacific | Papua New Guinea | 1978 | Cacao |
| P6298 | E196 | Asia-Pacific | Indonesia | 1989 | Cacao |
| P6972 | E197 | Asia-Pacific | Malaysia | 1990 | Vanda |
| P7506 | E198 | Asia-Pacific | Malaysia | 1991 | Durian |
| P0395 | E205 | America | Costa Rica | 1966 | Cacao |
| P0497 | E206 | America | Colombia | 1967 | Cacao |

**Supplementary Table 1** Continued

| Accession Number | DNA Number | Region | Country | Year | Host |
|---|---|---|---|---|---|
| P3548 | E207 | Asia-Pacific | Sri Lanka | - | Cacao |
| P6029 | E208 | America | Trinidad | 1989 | Cacao |
| P6284 | E209 | Asia-Pacific | Indonesia | 1989 | Coconut |
| P6282 | E210 | Asia-Pacific | Indonesia | 1989 | Coconut |
| P7548 | E211 | Asia-Pacific | Philippines | 1992 | Papaya |
| P8013 | E212 | Asia-Pacific | Australia | - | Baronia sp |
| P8030 | E213 | Asia-Pacific | Indonesia | - | Coconut |
| P8031 | E214 | Asia-Pacific | Indonesia | - | Coconut |
| P6595 | E215 | America | Florida, USA | 1989 | Bamboo Palm |
| P6966 | E216 | Asia-Pacific | Malaysia | 1990 | Cacao |
| P1915 | E217 | Asia-Pacific | French Polynesia | 1985 | Vanilla |
| P7541 | E220 | Asia-Pacific | Philippines | 1992 | Calamandarin |
| P8487 | E226 | Asia-Pacific | Sri Lanka | - | Rubber |

**Supplementary Table 2.** Primers designed for multilocus sequence analysis of *P. palmivora*.

| Locus | Primer name | Primer sequence |
|---|---|---|
| *β-tubulin* | btub_PpF1 | GTCCGAATTCTCCTCAGAGC |
| | btub_PpF1 | GTCCGAATTCTCCTCAGAGC |
| *trp1* | trp1_PpF1 | AAAGCGTTCGTCAGAAGCAT |
| | trp1_PpR1 | CGAGTCACGTTCTGCAAGAG |
| *pelota* | Pelota_PpF1 | GACTCGGTCAAGACGACGAC |
| | Pelota_PpR1 | CACGGTCCTGGTCAATATGC |

**Supplementary Table 3.** Nucleotide substitution model for each locus inferred from jModelTest, MCMC parameter settings, and root states with the highest posterior probabilities for each locus.

| Locus | BIC | | Model Used | MCMC | Sampling | Result |
|---|---|---|---|---|---|---|
| | BioNJ | ML | (Estimated) | Iteration Times | Frequency | |
| *β-tubulin* | HKY+I+G | HKY+I+G | HKY+I+G | $1.2 \times 10^9$ | $6.0 \times 10^4$ | Philippines and Indonesia |
| *pelota* | TPM2+I | TPM2+I | TPM2+I | $8.0 \times 10^8$ | $4.0 \times 10^4$ | Indonesia |
| *trp1* | HKY+I | HKY+I | HKY+I | $4.0 \times 10^8$ | $2.0 \times 10^4$ | Indonesia and Philippines |
| *coxII* and spacer | HKY+I | TrN+I | TrN+I | $8.0 \times 10^8$ | $4.0 \times 10^4$ | Philippines |

**Supplementary Table 4.** Inferred root state using three different starting seeds for the nuclear loci (a) *β-tubulin*, (b) *pelota*, (c) *trp1*, and (d) the mitochondrial *coxII* and spacer.

**a.** *β-tubulin*

| **Seed 1** | | **Seed 2** | | **Seed 3** | |
|---|---|---|---|---|---|
| **Root location** | **Prob.** | **Root location** | **Prob.** | **Root location** | **Prob.** |
| Philippines | 0.26 | Philippines | 0.26 | Philippines | 0.25 |
| Indonesia | 0.19 | Indonesia | 0.20 | Indonesia | 0.20 |

**b.** *pelota*

| **Seed 1** | | **Seed 2** | | **Seed 3** | |
|---|---|---|---|---|---|
| **Root location** | **Prob.** | **Root location** | **Prob.** | **Root location** | **Prob.** |
| Indonesia | 0.94 | Indonesia | 0.95 | Indonesia | 0.95 |

**c.** *trp1*

| **Seed 1** | | **Seed 2** | | **Seed 3** | |
|---|---|---|---|---|---|
| **Root location** | **Prob.** | **Root location** | **Prob.** | **Root location** | **Prob.** |
| Philippines | 0.50 | Philippines | 0.49 | Philippines | 0.48 |
| Indonesia | 0.50 | Indonesia | 0.51 | Indonesia | 0.51 |

**d.** *coxII and spacer*

| **Seed 1** | | **Seed 2** | | **Seed 3** | |
|---|---|---|---|---|---|
| **Root location** | **Prob.** | **Root location** | **Prob.** | **Root location** | **Prob.** |

| | | | | | |
|---|---|---|---|---|---|
| Philippines | 0.85 | Philippines | 0.73 | Philippines | 0.73 |

**Supplementary Table 5.** BASTA estimates of migration rates between two major hosts.

**a**. Two-host model

| | | Mean estimated immigration rate [95% HPD]: | |
|---|---|---|---|
| Source host | Root state probability | Cacao* | Coconut* |
| Cacao | 0.96-0.97 | - | **0.390 [0.099-0.740]** |
| Coconut | <0.1 | 0.056 [0-0.158] | - |

- Bold: lower boundary of 95% HPD > 0.001

**b**. Three-host model, including an unsampled host

| | | Mean estimated immigration rate [95% HPD]: | | |
|---|---|---|---|---|
| Source host | Root state probability | Cacao | Coconut | Unsampled |
| Cacao | 0.45-0.57 | - | **0.378 [0.086-0.734]** | 0.084 [0-0.254] |
| Coconut | <0.1 | 0.051 [0-0.149] | - | 0.086 [0-0.264] |

| | | | | |
|---|---|---|---|---|
| Unsampled | 0.41-0.51 | 0.130 [0-0.323] | 0.259 [0-0.577] | - |

Bold: lower boundary of 95% HPD > 0.001

**Supplementary Table 6.** BASTA estimates of migration rates between demes defined by host and geographical region. Models are those shown in Table 3 plus the Cacao-Africa deme.

**a**. Four-deme model*

| | | Mean estimated immigration rate [95% HPD]: | | | |
|---|---|---|---|---|---|
| Source deme | Root state probability | Cacao-America | Cacao-AsiaPacific | Cacao-Africa | Coconut-Asia |
| Cacao-America | 0.89-0.91 | - | **0.303 [0.054-0.573]** | **0.594 [0.188-1.052]** | 0.050 [0-0.138] |
| Cacao-AsiaPacific | <0.1 | 0.054 [0-0.151] | - | 0.072 [0-0.215] | **0.241 [0.053-0.465]** |
| Cacao-Africa | <0.1 | 0.089 [0-0.249] | 0.136 [0-0.382] | - | 0.144 [0-0.346] |
| Coconut-Asia | <0.1 | 0.024 [0-0.073] | 0.043 [0-0.129] | 0.038 [0-0.117] | - |

* *P.palmivora* isolates were not sampled from coconut in America; and Coconut-Africa deme is not included due to only one *P.palmivora* isolate in this deme.

Bold: lower boundary of 95% HPD > 0.001

**b**. Five-deme model, including an unsampled deme

| | | Mean estimated immigration rate [95% HPD]: | | | | |
|---|---|---|---|---|---|---|
| Source deme | Root state probability | Cacao-America | Cacao-AsiaPacific | Cacao-Africa | Coconut-Asia | Unsampled |
| Cacao-America | 0.68-0.73 | - | **0.335 [0.069-0.620]** | **0.626 [0.218-1.087]** | 0.055 [0-0.151] | 0.059 [0-0.183] |
| Cacao-AsiaPacific | <0.1 | 0.052 [0-0.150] | - | 0.075 [0-0.223] | **0.247 [0.034-0.490]** | 0.077 [0-0.232] |
| Cacao-Africa | <0.1 | 0.081 [0-0.230] | 0.132 [0-0.376] | - | 0.146 [0-0.354] | 0.077 [0-0.236] |
| Coconut-Asia | <0.1 | 0.027 [0-0.081] | 0.049 [0-0.145] | 0.042 [0-0.129] | - | 0.082 [0-0.244] |
| Unsampled | 0.19-0.22 | 0.066 [0-0.174] | 0.116 [0-0.329] | 0.076 [0-0.225] | 0.266 [0-0.592] | - |

Bold: lower boundary of 95% HPD > 0.001

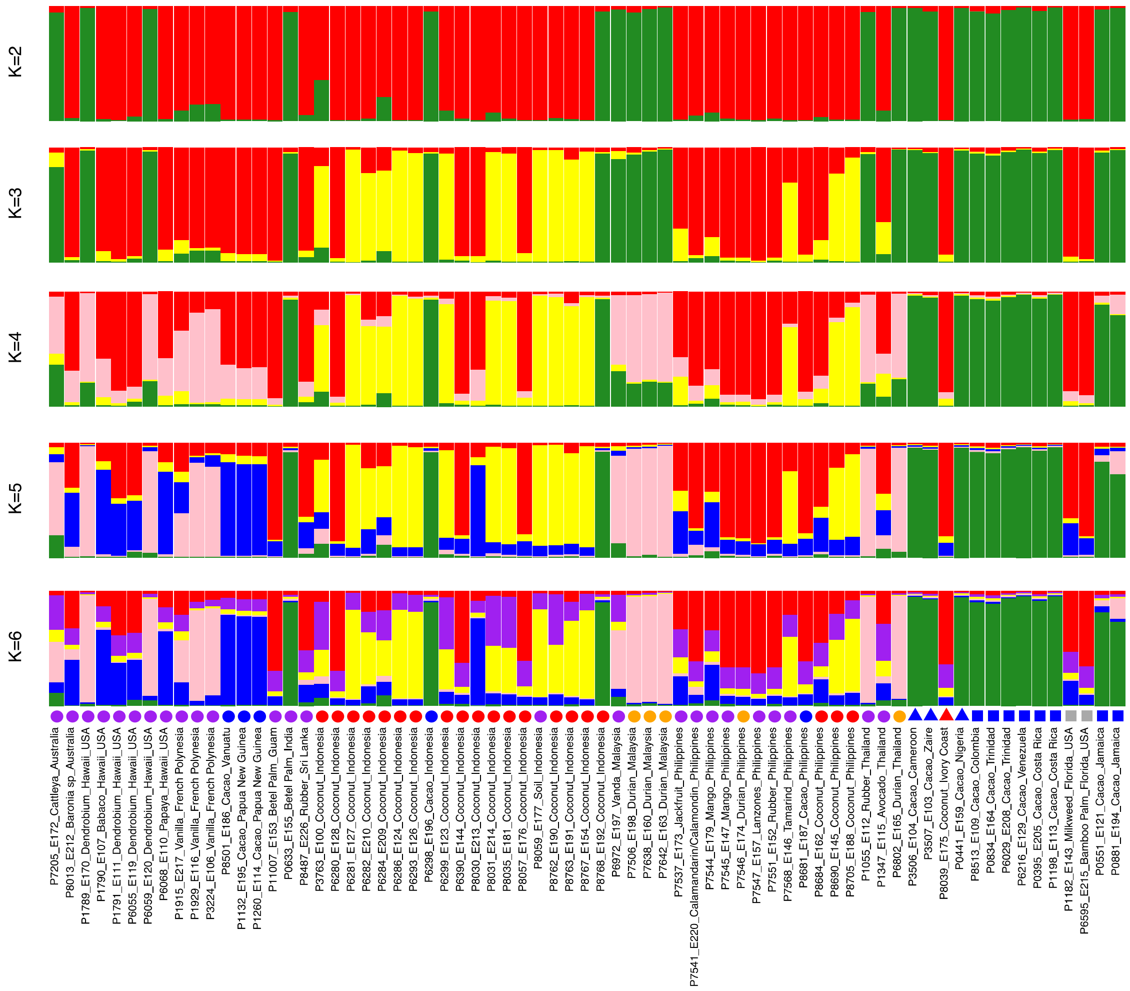


**Supplementary Figure 1. STRUCTURE assignments of clone-corrected *P. palmivora* isolates to populations for values of K from 2 to 6.** The optimum K value inferred from Evanno method was K=2. Greater genetic structure is still apparent in the Asia-Pacific region. The symbols represent the same meaning in the legend of Figure 2.

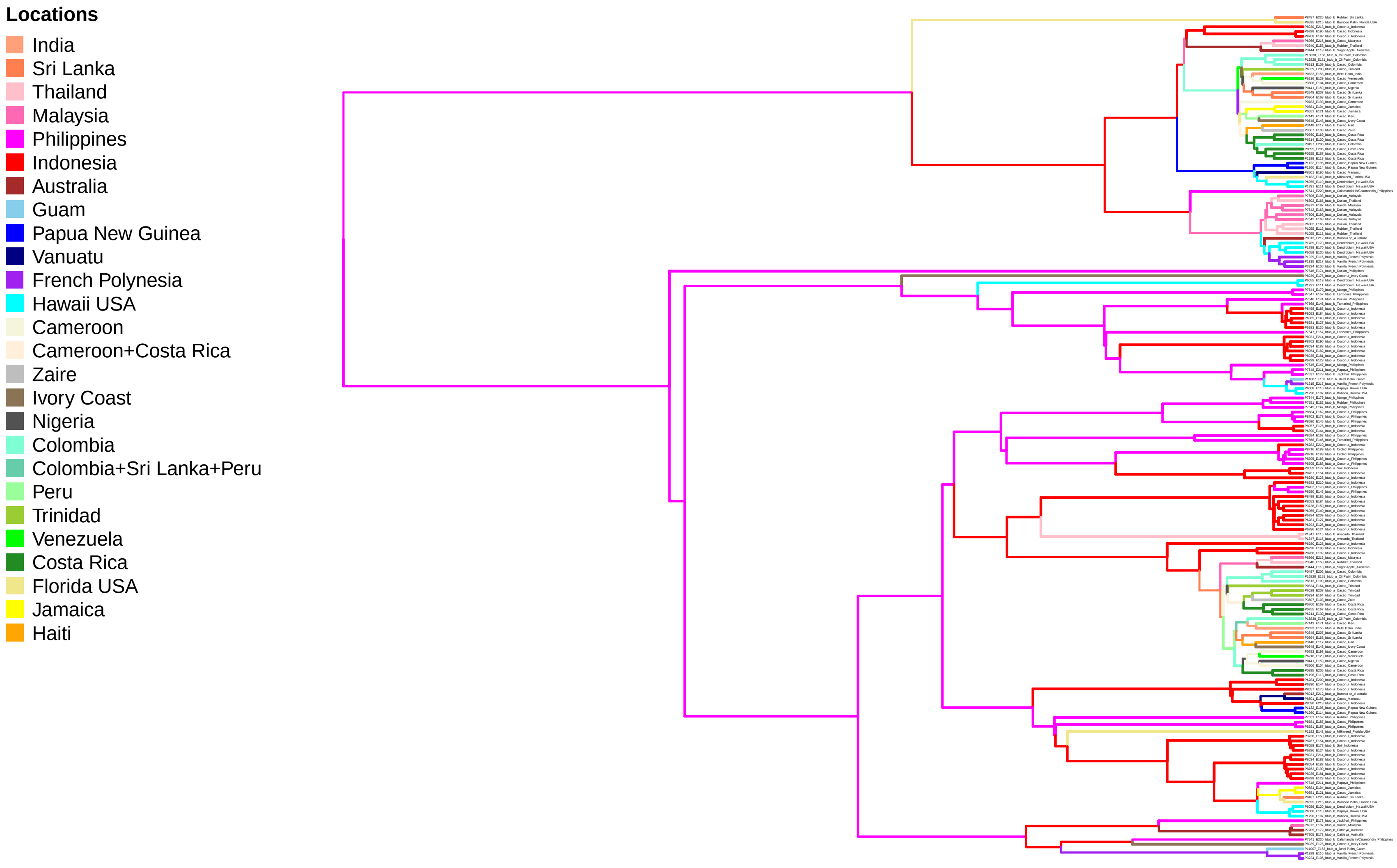


**Supplementary Figure 2. Maximum clade credibility genealogy for the *β-tubulin* locus.** Colors of branches indicate the most probable geographic origin of each lineage based on discrete phylogeographic analysis.

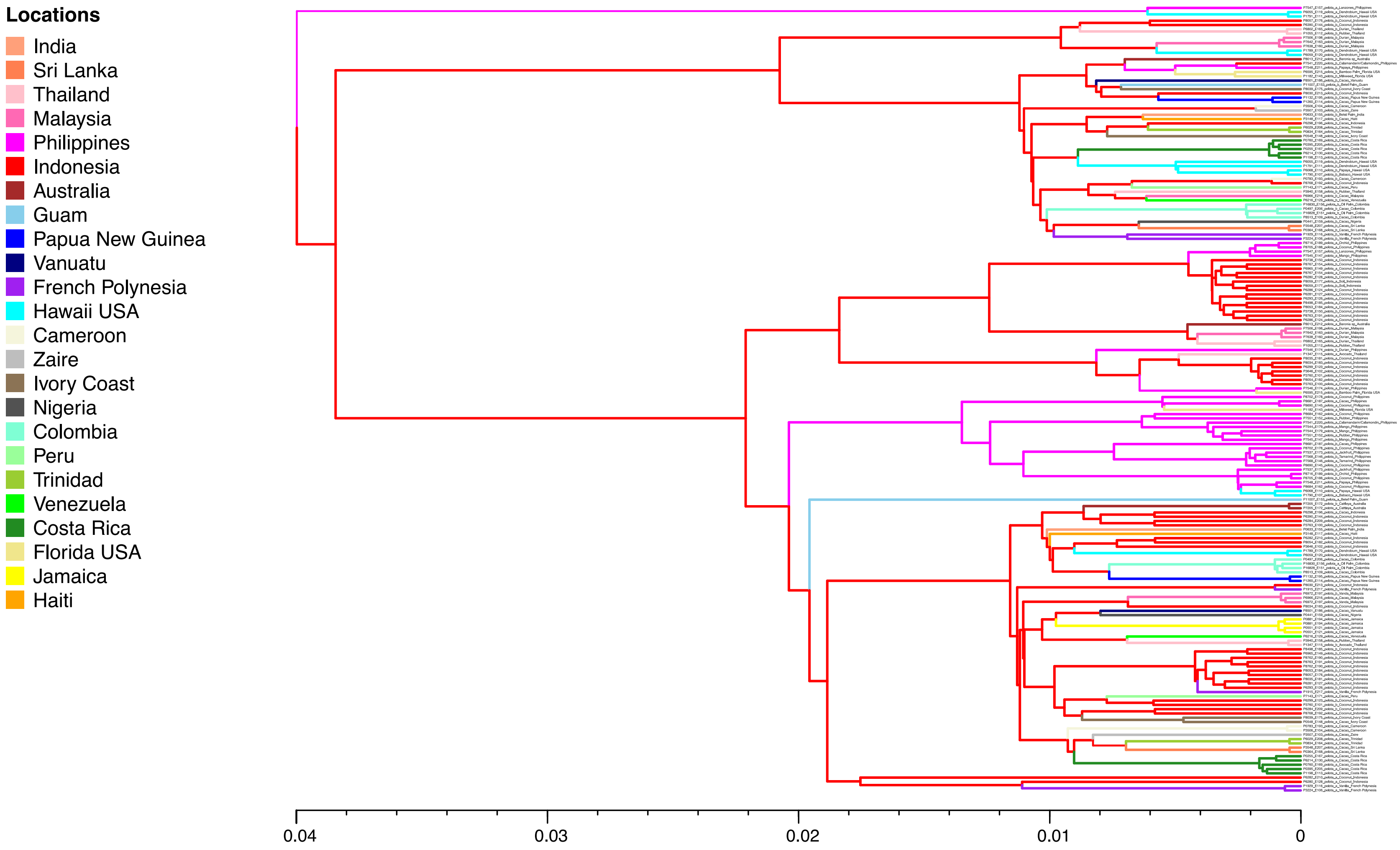


**Supplementary Figure 3. Maximum clade credibility genealogy for the *pelota* locus.** Colors of branches indicate the most probable geographic origin of each lineage based on discrete phylogeographic analysis.

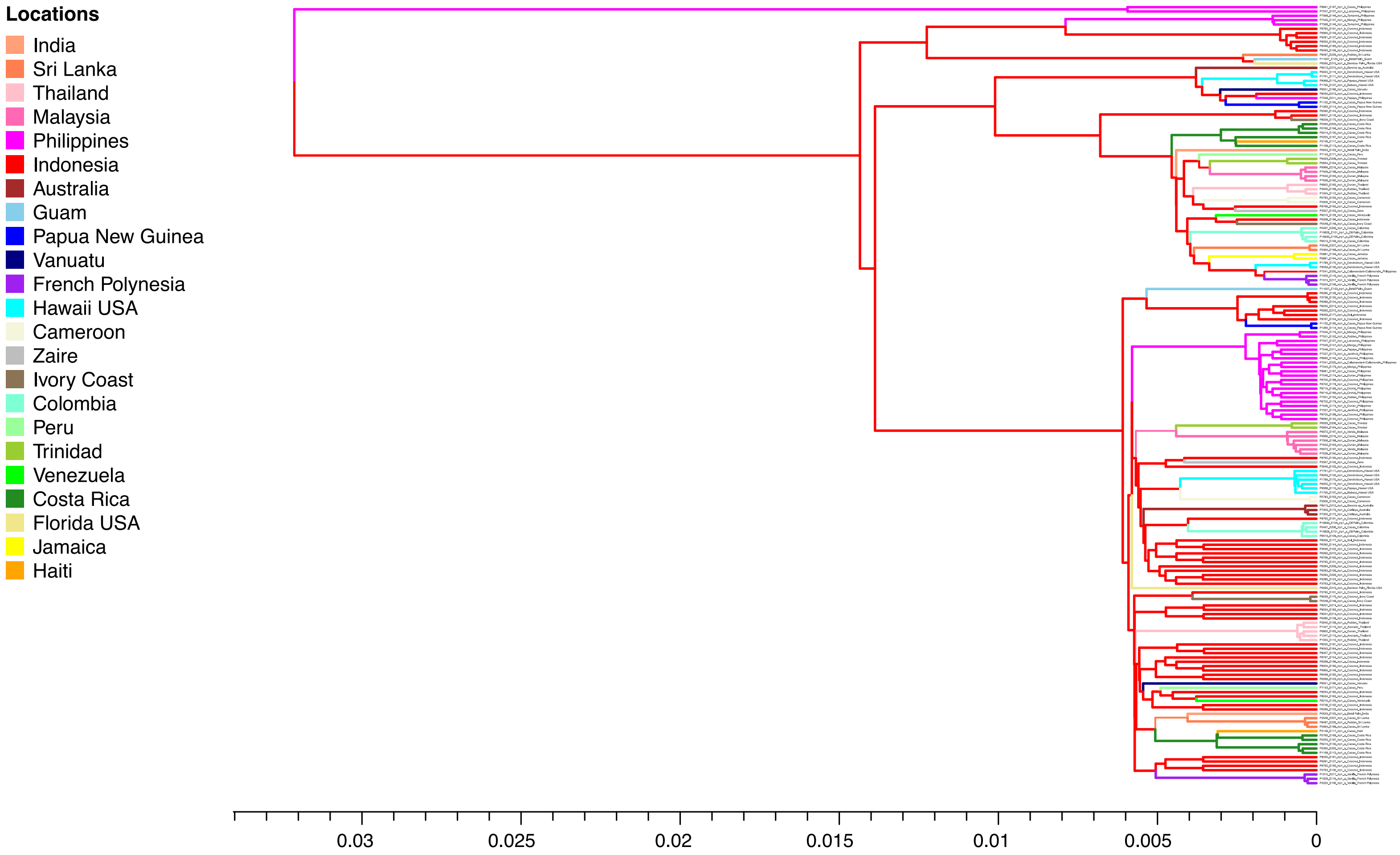


**Supplementary Figure 4. Maximum clade credibility genealogy for the *trp1* locus.** Colors of branches indicate the most probable geographic origin of each lineage based on discrete phylogeographic analysis.

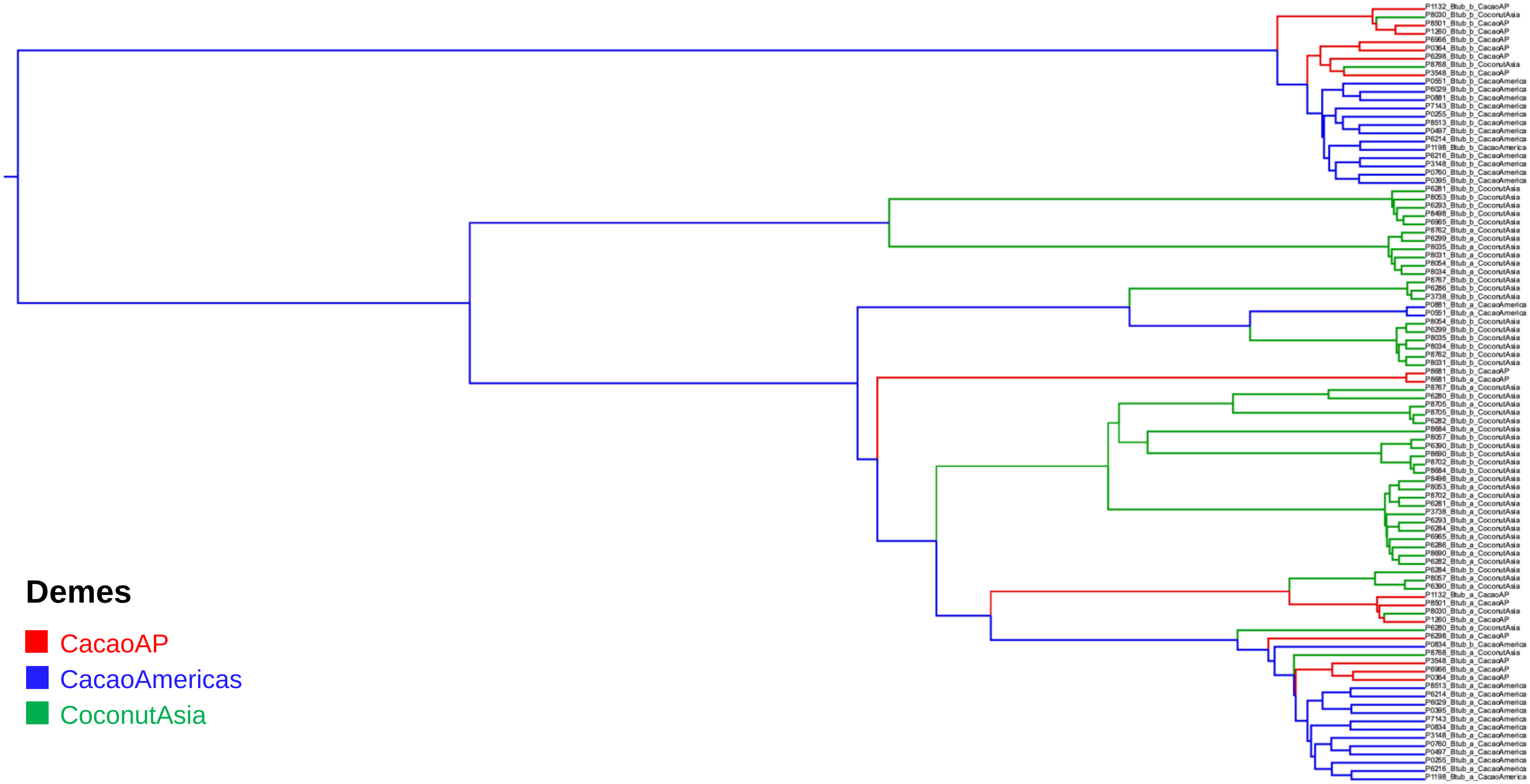


**Supplementary Figure 5. Maximum clade credibility genealogy for the *β-tubulin* locus using isolates collected from cacao and coconut in Asia-Pacific and the Americas.** Colors of branches indicate the most probable deme among the three demes: cacao in Asia-Pacific region, cacao in Americas, and coconut in Asia.

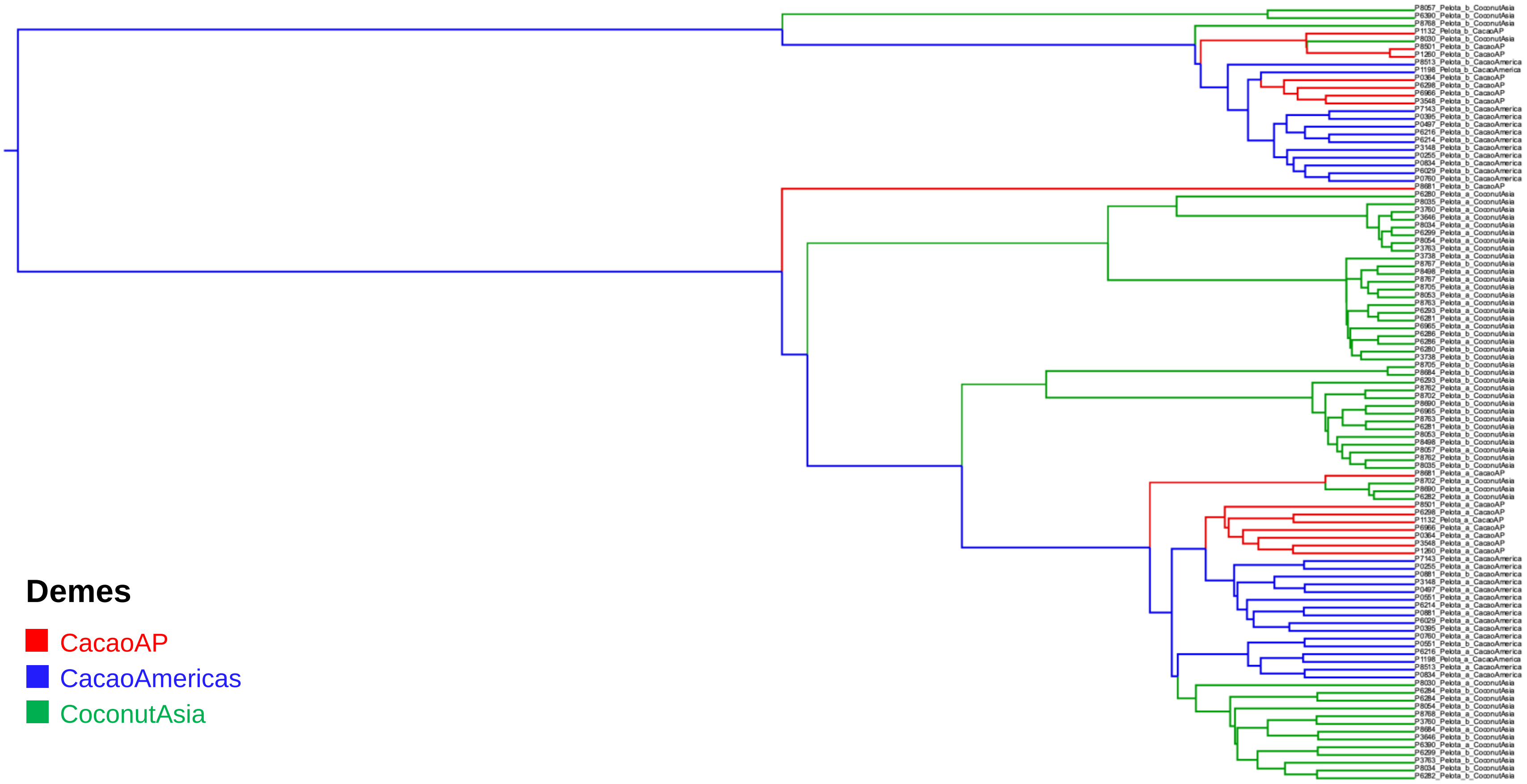


**Supplementary Figure 6. Maximum clade credibility genealogy for the *pelota* locus using isolates collected from cacao and coconut in Asia-Pacific and the Americas.** Colors of branches indicate the most probable deme among the three demes: cacao in Asia-Pacific region, cacao in Americas, and coconut in Asia.

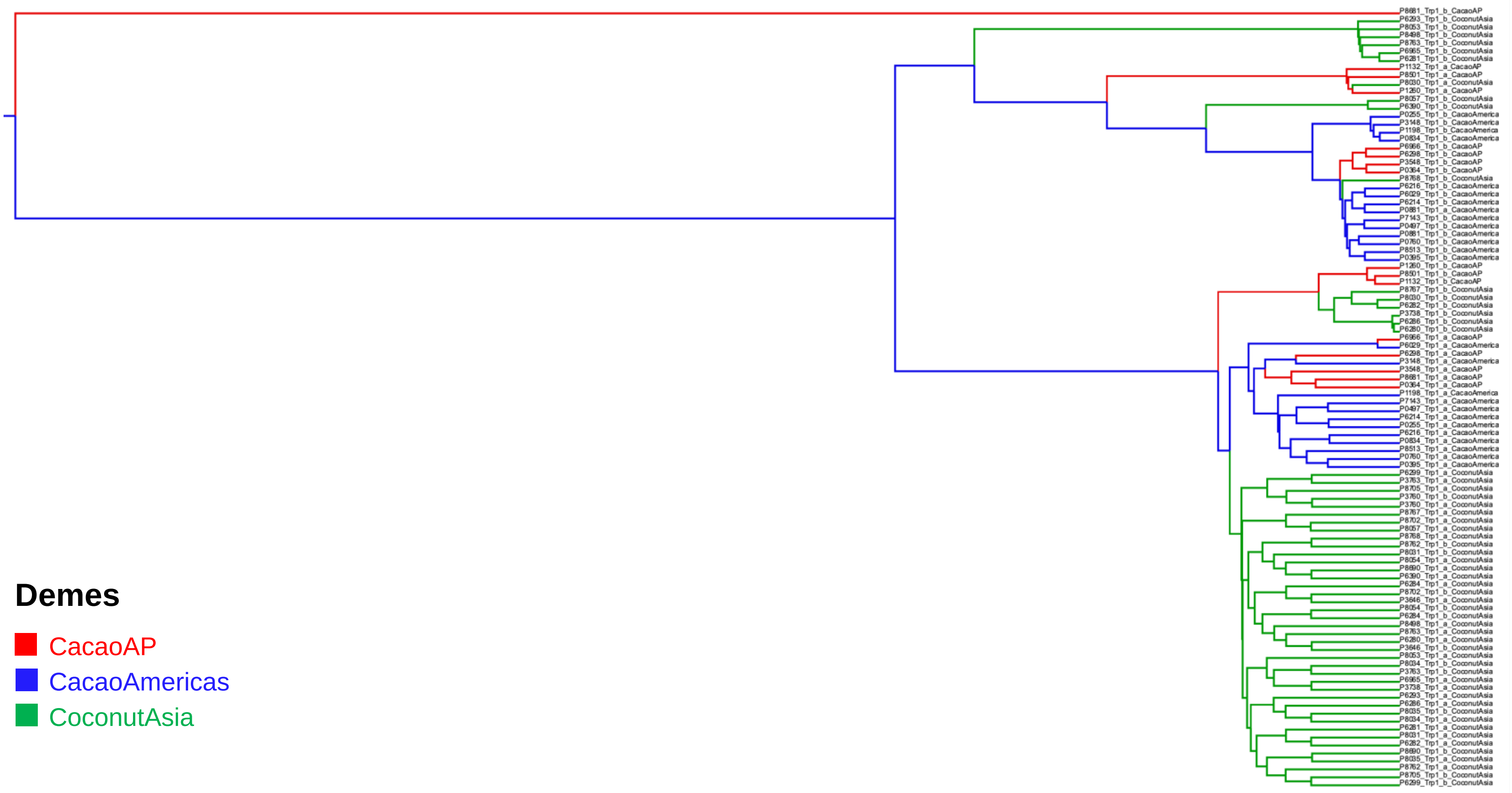


**Supplementary Figure 7. Maximum clade credibility genealogy for the *trp1* locus using isolates collected from cacao and coconut in Asia-Pacific and the Americas.** Colors of branches indicate the most probable deme among the three demes: cacao in Asia-Pacific region, cacao in Americas, and coconut in Asia.

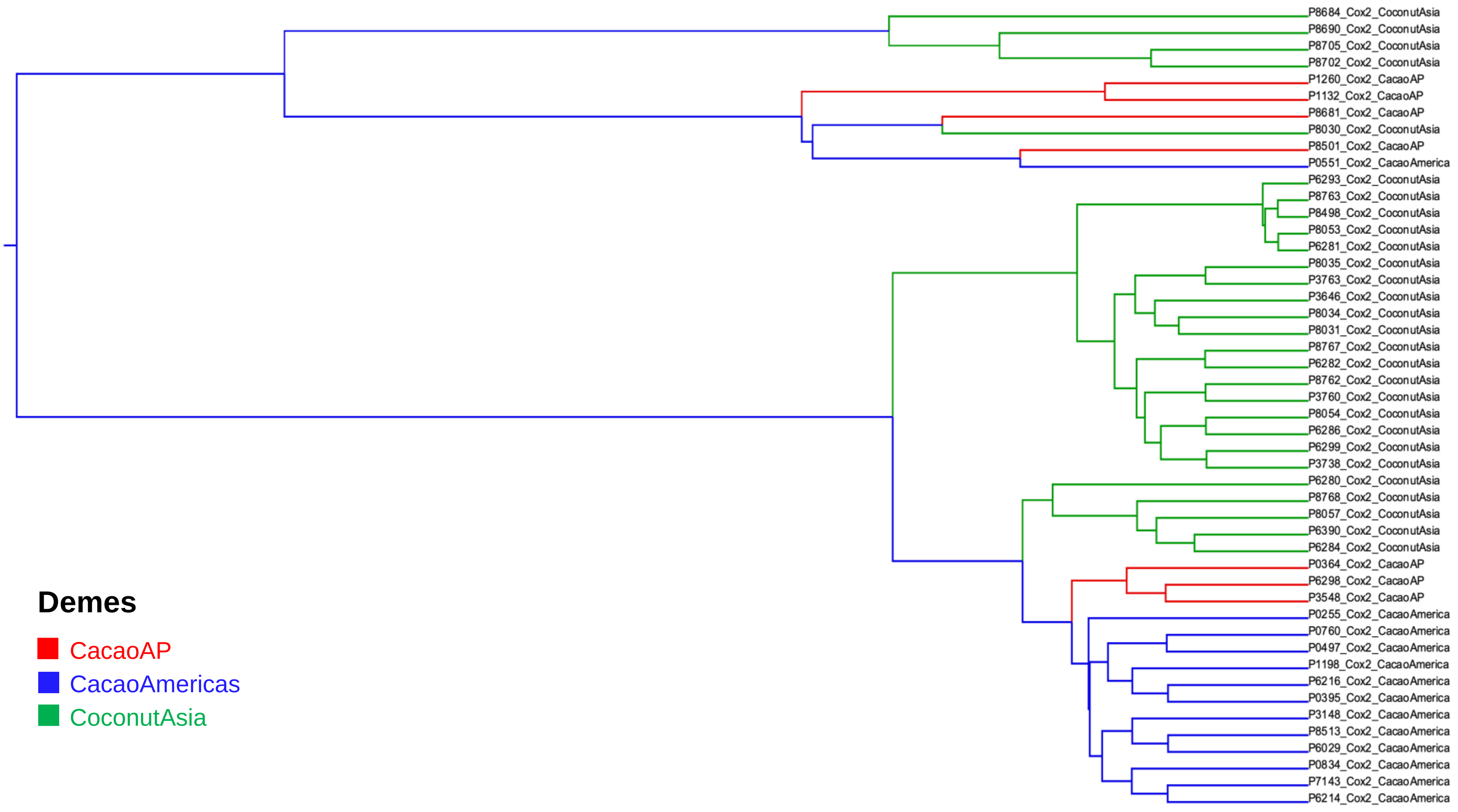


**Supplementary Figure 8. Maximum clade credibility genealogy for the *coxII* locus using isolates collected from cacao and coconut in Asia-Pacific and the Americas.** Colors of branches indicate the most probable deme among the three demes: cacao in Asia-Pacific region, cacao in Americas, and coconut in Asia.
